# Neurovascular Mechanisms of Cognitive Aging: Sex-Related Differences in the Average Progression of Arteriosclerosis, White Matter Atrophy, and Cognitive Decline

**DOI:** 10.1101/2023.09.06.556562

**Authors:** Daniel C. Bowie, Kathy A. Low, Samantha L. Rubenstein, Samia S. Islam, Benjamin Zimmerman, Paul B. Camacho, Bradley P. Sutton, Gabriele Gratton, Monica Fabiani

**Affiliations:** Department of Psychology, University of Illinois Urbana-Champaign, 603 E Daniel St., Champaign, IL 61820; Beckman Institute for Advanced Science and Technology, University of Illinois Urbana-Champaign, 405 N. Mathews Ave., Urbana, IL, 61801; Helfgott Research Institute, National University of Natural Medicine, 2220 SW 1st Ave., Portland, OR, 97201; Department of Bioengineering, University of Illinois Urbana-Champaign, 1406 W Green St, Urbana, IL, 61801

**Keywords:** cognitive aging, cerebrovascular health, cerebral arterial pulse based on diffused optical tomography (pulse-DOT), arteriosclerosis, white matter lesions, sex differences

## Abstract

Arterial stiffness (arteriosclerosis) has been linked to heightened risks for cognitive decline, and ultimately for Alzheimer’s disease and other forms of dementia. Importantly, neurovascular outcomes generally vary according to one’s biological sex. Here, capitalizing on a large sample of participants with neuroimaging and behavioral data (*N* = 203, age range = 18-87 years), we aimed to provide support for a hierarchical model of neurocognitive aging, which links age-related declines in cerebrovascular health to the rate of cognitive decline via a series of intervening variables, such as white matter integrity. By applying a novel piecewise regression approach to our cross-sectional sample to support Granger-like causality inferences, we show that, on average, a precipitous decline in cerebral arterial elasticity (measured with diffuse optical imaging of the cerebral arterial pulse; pulse-DOT) *temporally precedes* an acceleration in the development of white matter lesions by nearly a decade, with women protected from these deleterious effects until approximately age 50, the average onset of menopause. By employing multiple-mediator path analyses while controlling for sex, we show that age may impair cognition via the sequential indirect effects of arteriosclerosis and white matter atrophy on fluid, but not crystallized, abilities. Importantly, we replicate these results using pulse pressure, an independent index of arterial health, thereby providing converging evidence for the central role of arteriosclerosis as an accelerating factor in normal and pathological aging and identifying robust sex-related differences in the progression of cerebral arteriosclerosis and white matter degradation.

## 1. INTRODUCTION

### 1.1 Age-Related Declines in Cerebrovascular and White Matter Health

Cerebrovascular health declines during aging due, in part, to the gradual stiffening of cerebral arteries, or loss of arterial elasticity (arteriosclerosis), and consequent decreases in perfusion of oxygenated blood to neural tissue (Fabiani et al., 2022; Zimmerman et al., 2021). It is believed that vascular risk factors including hypertension and diabetes contribute to both an increased risk for Alzheimer’s disease (AD) pathology (e.g., increased amyloid beta burden) and non-AD cerebrovascular pathologies (e.g., white matter lesions and cerebral infarcts; Villeneuve & Jagust, 2015), albeit via largely independent pathways. Independent or not, the most common form of dementia remains mixed dementia, involving both AD and cerebrovascular pathologies (Schneider et al., 2007). Therefore, it is crucial to investigate the neurovascular mechanisms implicated in “normal” and “pathological” cognitive aging, thereby permitting identification of potential points for intervention.

The present study, utilizing a multimodal neuroimaging approach, provides multiple sources of converging evidence for the deleterious role that age-related declines in cerebrovascular health play in overall brain health and, ultimately, cognition., We also examined the possible differential effects of age on cerebrovascular health for men and women, given the vasoprotective role of estrogens in women until menopause (Reslan & Khalil, 2012; Xing et al., 2009).

It is widely known through large cross-sectional epidemiological studies that the first signs of arterial dysfunction become apparent in middle age (50-60 years old; Burt & Harris, 1994; Mahmood et al., 2014; Staessen et al., 2000; Stamler & Neaton, 2008). This leads to specific predictions regarding the *average* trajectories of phenomena that are likely consequences of arterial dysfunction. Namely, when examined with a novel piecewise regression approach applied to a cross-sectional adult sample, the group average trajectories should show minimal changes before the onset of arteriosclerosis, and variably lagged declines thereafter. This may offer important clues about their significance within the model cascade presented in **Figure 1**, allowing for Granger-like, but not direct, causal inferences. These effects should be less pronounced or delayed in women compared to men. It is believed that arteriosclerosis and atherosclerosis (the partial blockage or narrowing of arteries due to cholesterol plaque accumulation) ultimately result in partial ischemia and contribute to chronic cerebral hypoperfusion (Li et al., 2018) and to the development of cerebral small vessel disease (CSVD; see Chojdak-Łukasiewicz et al., 2021 for a review). CSVD, a chronic and progressive neurovascular disease, primarily affects smaller cerebral blood vessels (< 400 µm in diameter) such as the smaller arteries and arterioles that supply blood to the brain’s white matter and subcortical gray matter. In its acute form, CSVD accounts for 25% of all strokes, including 25% of all ischemic strokes. However, even in the absence of a symptomatic cerebrovascular accident, the presence of CSVD can be estimated on the basis of hallmark neuroimaging findings, including white matter signal abnormalities (WMSAs; hypointense regions on T1-weighted MRI images or hyperintense regions on T2-FLAIR MRI images; Pantoni et al., 1996; Pantoni & Garcia, 1997). WMSAs are non-specific lesions and may indicate degeneration of the myelin sheath due to the loss of myelin-forming oligodendrocytes, axonal loss, or reactive gliosis (Pantoni & Garcia, 1997). The loss of electrically insulating myelin around neuronal axons decreases the velocity at which action potentials are conducted, limiting rates of neuronal firing and information exchange between neurons.

**Figure 1:**
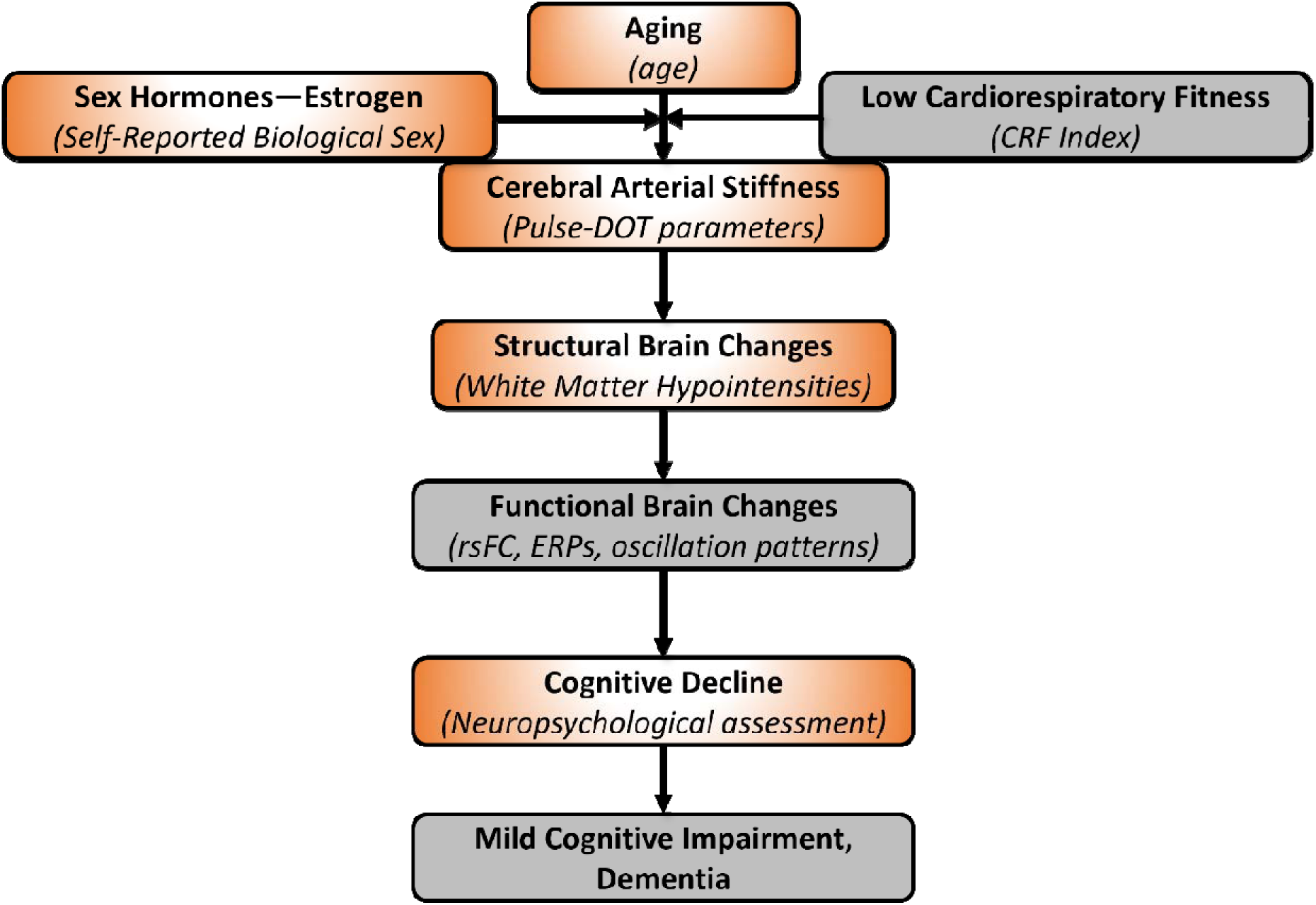
Hypothesized arterial dysfunction pathway to cognitive decline (measures). Evidence suggests that CRF may moderate the relationship between age and arterial health, thereby providing a point of intervention (via regular aerobic exercise) to prevent or slow the age-related progression of arteriosclerosis. Female sex hormones such estradiol may also confer protective effects to the cerebrovasculature prior to menopausal onset, after which production of estrogens declines precipitiously around age 50. Greyed out panels indicate variables for which we do not have measures of in the present paper. CRF = cardiorespiratory fitness; pulse-DOT = diffuse optical tomography of the cerebral arterial pulse; rsFC = resting-state fMRI functional connectivity; ERPs = event-related brain potentials; MCI = mild cognitive impairment

Cerebrovascular health can be estimated noninvasively by examining indices of cerebral blood flow derived from magnetic resonance imaging (e.g., arterial spin labeling; Calamante et al., 1999) or Transcranial Doppler ultrasound (Purkayastha & Sorond, 2012). Near-infrared spectroscopy (NIRS), an optical imaging technique, can also provide noninvasive estimates of cerebrovascular health based on analyses of the cerebral pulse waveform, which represents oxygenation-dependent changes in near-infrared light absorption. Prior research utilizing this technique, known as diffuse optical tomography of the cerebral arterial pulse (pulse-DOT), has demonstrated that various parameters of the optical cerebral pulse are reliably associated with age, cardiorespiratory fitness, brain tissue atrophy (globally and regionally), and brain function (Chiarelli et al., 2017; Fabiani et al., 2014; Gratton et al., 2017; Kong et al., 2020; Tan et al., 2017, 2019). By examining the optical pulse waveform time-locked to the electrocardiogram (EKG) R wave (indicating peak ventricular depolarization), one can extract several indices of cerebrovascular health, including, but not limited to, the pulse relaxation function (PReFx; Fabiani et al., 2014). PReFx describes the temporal overlap between the forward and backward pressure waves generated during each cardiac cycle. A greater overlap between these two waves is associated with low PReFx values and indexes arteriosclerosis, whereas a small overlap is associated with high PReFx values and higher arterial elasticity. Also, importantly, PReFx refers to *cerebral* arterial elasticity in the arterial tract connecting the point of measurement with the place where peripheral resistance occurs, and the backward (reflected) pressure wave is generated. This wave’s formation occurs downstream from the measurement point. Therefore, despite being superficially measured, PReFx exhibits sensitivity not only to the arterial stiffening occurring in the cortical mantle but also to stiffness occurring in deeper regions, including those prone to white matter abnormalities. The present paper utilizes PReFx as the primary cerebral index of neurovascular health.

Based on prior evidence, Kong and colleagues (2020) proposed a hierarchical model of neurocognitive aging that positions impaired cerebrovascular integrity as one of the early cascading factors that detrimentally impact brain structure, function, and ultimately cognitive well-being (see **Figure 1** for an updated diagram of this mechanistic model). While this model is not exhaustive or fully representative of all possible causal relations or pathways to cognitive decline, it emphasizes the role of cerebrovascular health, whose decline with age can be mitigated with regular aerobic exercise, thereby providing a potential target for interventions to prevent or slow down age-related neurocognitive decline (Colcombe & Kramer, 2003; Hillman et al., 2008). Although their findings were generally consistent with the proposed model, the cross-sectional correlational analysis and modest sample size (*N* = 48) precluded strong inferences regarding the postulated causal role of cerebrovascular mechanisms in cognitive aging.

### 1.2 Hypotheses of Present Study

The present study expands upon the preliminary findings of Kong et al. (2020) by increasing the sample size more than four-fold (*N* = 203), conducting piecewise regression and multiple-mediator path analyses (i.e., mediation analyses involving multiple mediators/intervening variables in succession), and examining pertinent sex-related differences, thereby enabling group-level Granger-like causal inferences. To accomplish this, we pooled and harmonized data from several in-lab studies (see section 2.3 and Supplementary Materials), in which key measures of interest were obtained, including participants’ age, cerebral arterial elasticity (pulse-DOT PReFx; Fabiani et al., 2014; Tan et al., 2017; Gratton et al., 2017), neuroanatomical integrity (WMSA volume), and fluid cognition (a composite measure based on multiple neuropsychological tests). In fact, fluid cognition is particularly susceptible to age-related decline, compared to crystallized cognition, which grows to adulthood and is maintained until old age (Salthouse, 2012). We also sought to replicate the results of all analyses by replacing PReFx, an index of cerebrovascular health, with a systemic index of arterial health, pulse pressure (Stanley S Franklin, 2006; Izzo & Shykoff, 2001; Steppan et al., 2011). Pulse pressure is the difference between systolic and diastolic blood pressure, which is inversely related to arterial compliance and generally increases with age, starting in middle age. These additional analyses provide an external validation for the pulse-DOT approach.

The piecewise (or segmented) regressions allowed us to model the directional relationships between age and arteriosclerosis (as indexed by PReFx and pulse pressure), age and WMSAs (T1-weighted white matter hypointensity volumes), age and fluid cognition, and age and crystallized cognition, which are hypothesized in **Figure 1**. Specifically, this analytic approach allows us to identify “breakpoints” or “knees” joining two linear regression functions of significantly different slopes. That is, this allows us to identify the approximate age ranges during which declines in cerebral arterial elasticity, white matter health, and fluid or crystallized cognition begin or accelerate at the group level, overall and by sex. Of course, the employment of a cross-sectional design, as always, should caution readers against drawing unwarranted conclusions, as confounding cohort effects are ever-present. Nonetheless, given the difficulties, time duration, and costs inherent to performing long-term longitudinal studies, this approach allows us to model these complex relationships in a cross-sectional sample and may offer additional insights into the general neurovascular mechanisms implicated in cognitive aging and important sex-related differences.

## 2. METHOD

### 2.1 Participants

Neuroimaging and behavioral data from four lab studies were pooled and harmonized to increase statistical power for a cross-sectional investigation of some of the relationships amongst variables in the hypothesized cascade model of neurocognitive aging depicted in **Figure 1**. *Despite the use of pooled data, please note that all the analyses presented here are previously unpublished*. The pooled sample included ***N* = 203 unique participants** (see **Table 1** for sample size, self-reported sex and age-related statistics sorted by study).

**Table 1:**
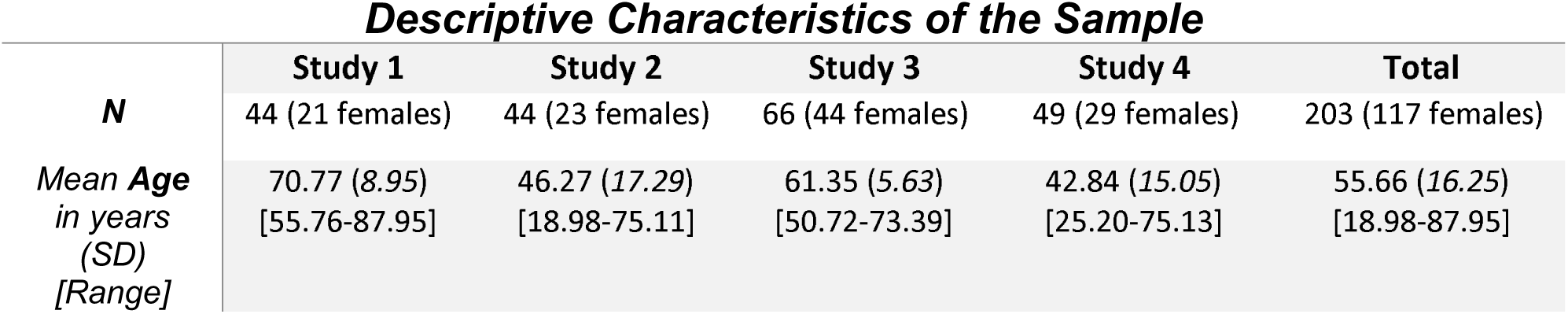
Descriptive characteristics of the sample arranged by study. The rightmost column includes statistics for all participants across all studies.

Participants in studies 1 and 2 reported no history of neurological or psychiatric disorders and had no signs of dementia (as assessed by a score > 51 on the modified Mini-Mental Status Examination (MMSE; Mayeux et al., 1981)) or depression (as assessed by the Beck’s Depression Inventory; (Beck et al., 1996)). Participants in studies 3 and 4 were administered the Montreal Cognitive Assessment and were included if they scored > 20 points^1^ Nasreddine et al. (2005) suggest that cognitively intact participants score 26-30 points, while participants scoring 18-25 possibly suffer from mild cognitive impairment (MCI). Therefore, studies 3 and 4 may include some participants who may have suffered from MCI during assessments and neuroimaging sessions, although variability in SES and education can also affect the score range. (Nasreddine et al., 2005). Informed consent was provided by each participant. All procedures were approved by the Institutional Review Board of the University of Illinois.

### 2.2 Data Acquisition

#### 2.2.1 Neuropsychological Assessment

Across all four studies, participants were administered an array of neuropsychological tests, including the Trail Making Tests A and B (Corrigan & Hinkeldey, 1987) to assess processing speed and working memory, the OSPAN task (Unsworth et al., 2005) to assess working memory capacity, and other tests of fluid and crystallized abilities. Given that different, non-overlapping tests of the same cognitive construct were sometimes administered across studies (e.g., Wisconsin Card Sorting Test vs. Dimensional Change Card Sort Test—NIH Toolbox, both of which assess cognitive flexibility), statistical harmonization of these data was required to ensure that scores were comparable. We employed a procedure that allowed us to compute standardized scores based on a multitude of tests (some of which overlapped and some of which did not) assessing crystallized ability and fluid abilities. This procedure is summarized in section 2.3.1, with a more detailed explanation and validation of the procedure in **supplementary materials**. The tests administered and statistically harmonized are listed in **Table 2**.

**Table 2:**
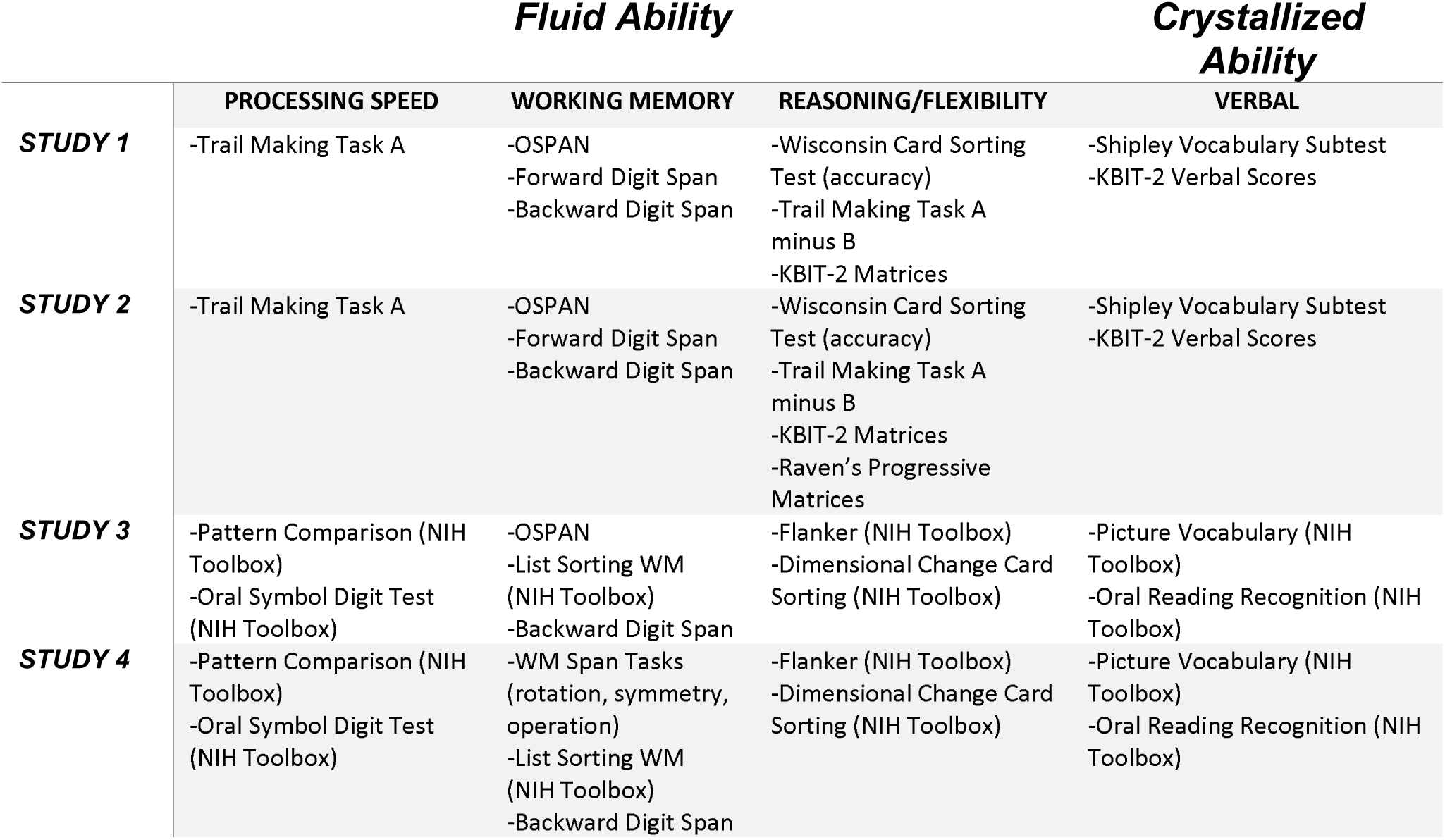
Neuropsychological tests (sorted by study) administered to assess various cognitive constructs (sorted by column). All assessments under the ‘Fluid Ability’ heading (processing speed, working memory, reasoning/flexibility) were statistically harmonized and used to compute a composite fluid ability score. The same was done for tests under the ‘Crystallized Ability’ heading (verbal), comprising tests that assess crystallized ability.

|  | ***Fluid Ability*** |  |  | ***Crystallized Ability*** |
| --- | --- | --- | --- | --- |
|  | **PROCESSING SPEED** | **WORKING MEMORY** | **REASONING/FLEXIBILITY** | **VERBAL** |
| **STUDY 1** | -Trail Making Task A | -OSPAN<br>-Forward Digit Span<br>-Backward Digit Span | -Wisconsin Card Sorting Test (accuracy)<br>-Trail Making Task A minus B<br>-KBIT-2 Matrices | -Shipley Vocabulary Subtest<br>-KBIT-2 Verbal Scores |
| **STUDY 2** | -Trail Making Task A | -OSPAN<br>-Forward Digit Span<br>-Backward Digit Span | -Wisconsin Card Sorting Test (accuracy)<br>-Trail Making Task A minus B<br>-KBIT-2 Matrices<br>-Raven's Progressive Matrices | -Shipley Vocabulary Subtest<br>-KBIT-2 Verbal Scores |
| **STUDY 3** | -Pattern Comparison (NIH Toolbox)<br>-Oral Symbol Digit Test (NIH Toolbox) | -OSPAN<br>-List Sorting WM (NIH Toolbox)<br>-Backward Digit Span | -Flanker (NIH Toolbox)<br>-Dimensional Change Card Sorting (NIH Toolbox) | -Picture Vocabulary (NIH Toolbox)<br>-Oral Reading Recognition (NIH Toolbox) |
| **STUDY 4** | -Pattern Comparison (NIH Toolbox)<br>-Oral Symbol Digit Test (NIH Toolbox) | -WM Span Tasks (rotation, symmetry, operation)<br>-List Sorting WM (NIH Toolbox)<br>-Backward Digit Span | -Flanker (NIH Toolbox)<br>-Dimensional Change Card Sorting (NIH Toolbox) | -Picture Vocabulary (NIH Toolbox)<br>-Oral Reading Recognition (NIH Toolbox) |

#### 2.2.2 Structural MRI Acquisition

For participants in studies 1 and 2 identical high-resolution structural scans were collected in a 3-Tesla Siemens Trio MR scanner, using a 12-channel head coil. The 3D T1-weighted anatomical scan for each participant was acquired using an MPRAGE sequence with the following parameters: TR = 1,900 ms; TE = 2.32 ms; TI = 900 ms; 192 sagittal slices; slice thickness = 0.90 mm; FA = 9°; voxel sizes = 0.9 × 0.9 × 0.9 mm, field-of-view (FOV) = 172.8 × 230 × 230 mm.

Participants’ T1-weighted images in studies 3 and 4 were acquired in a 3-Tesla Siemens Prisma MR scanner, using a 20-channel head coil. The MPRAGE sequence utilized the following pulse parameters: TR = 2,400 ms; TE = 2.31 ms; TI = 1,060 ms; 224 sagittal slices; slice thickness = 0.80 mm; FA = 8°; voxel sizes = 0.8 × 0.8 × 0.8 mm, field-of-view (FOV) = 179.2 × 256 × 256 mm, acceleration factor = 2.

#### 2.2.3 Diffuse Optical Imaging data acquisition

Cerebral arterial elasticity (pulse-DOT) data were obtained during a resting-state optical imaging session, in which seated participants fixated on a cross in the center of a screen. Optical data were acquired with a multichannel frequency-domain oximeter (ISS Imagent, Champaign, Illinois) equipped with 128 laser diodes (64 emitting light at 690 nm and 64 at 830 nm) and 24 photomultiplier tubes. Time-division multiplexing was employed so that each detector picked up light from 16 different sources at different times within a multiplexing cycle at a sampling rate of 39.0625 Hz. The light was transmitted to the scalp by using single-optic fibers (0.4-mm core) and from the scalp back to the photomultiplier tubes by using fiber bundles (3-mm diameter). The fibers were held in place using comfortable semirigid custom-built helmets, fitted to participants based on their head circumference.

After the helmet was set up, the locations of the optodes were marked digitally to improve spatial accuracy during later data processing, including anatomical coregistration with each participant’s structural MRI. Fiducial markers were placed on each participant’s left and right preauricular points and on the nasion. These fiducial points, optode locations, and other scalp locations were digitized with a Polhemus FastTrak 3D digitizer (accuracy: 0.8 mm; Colchester, VT) by using a recording stylus and three head-mounted receivers, which allowed for small movements of the head in between measurements. Optode locations and structural MRI data were then coregistered using the fiducials and a surface-fitting Levenberg and Marquardt algorithm (Chiarelli et al., 2015).

Concurrently, lead 1 of the EKG (left wrist referenced to right wrist) was recorded using a Brain-Vision recorder and a Brain Products V-Amp 16 integrated amplifier system (Brain Products, Germany), with a sampling rate of 500 Hz and a band-pass filter of 0.1–100 Hz. The exact timing of each R-wave peak was determined by searching for peak points exceeding a voltage threshold (scaled for each participant) and dismissing any peak points outside the normal range of inter-beat intervals. The identification of each peak was verified by visual inspection, and false detections, usually misidentifying the T-wave as the R-wave, were eliminated. Concurrent acquisition of EKG and optical pulse data allowed us to time-lock the optical pulse data to the R wave of the EKG, thereby ensuring that the same pulse was analyzed irrespective of its location within the brain. Optical pulse data were acquired from most of the scalp surface but with varying degrees of density (indexed by the number of channels) depending on the specific study. Optical data from study 1 were acquired via one montage (i.e., a single source-detector configuration) comprising 384 channels. Study 2 utilized four montages, each containing 384 channels, for a total of 1,536 channels. Study 3 utilized two montages, each containing 768 channels, for a total of 1,536 channels. Lastly, data for study 4 were acquired for one montage containing 768 channels.

#### 2.2.4 Systemic Pulse Pressure

In all four studies, participants’ blood pressure was taken at the brachial artery of the upper arm at three time points across the experiment and averaged to provide reliable measures of systolic and diastolic blood pressure. Systemic pulse pressure was computed by taking the difference between systolic and diastolic blood pressure. This served as a methodologically independent index of systemic arterial elasticity that provided us the opportunity to validate the results observed when using pulse-DOT as an index of cerebrovascular health. All plots and mediation diagrams involving pulse pressure are presented in the **supplementary materials,** but related results are briefly summarized in the main text when necessary.

### 2.3 Data Processing and Statistical Harmonization

For all variables (except age), values greater or less than three standard deviations from the sample mean were excluded as outliers. Subsequently, WMSA volumes (here, T1-weighted white matter hypointensity volumes), which were positively skewed, were log-transformed to be approximately normal.

#### 2.3.1 Neuropsychological Data

To statistically harmonize measures tapping into overlapping cognitive constructs across studies, we computed standardized scores of crystallized and fluid abilities for each participant. Fluid ability consists of scores on tests assessing processing speed, working memory, cognitive flexibility, and reasoning, which are highly vulnerable to age-related neurobiological decline (Hedden & Gabrieli, 2004; see **Table 2** for the complete list of tests used to compute composite fluid and crystallized ability scores). Please view the **supplementary materials** for a more detailed description and validation of the procedure, in which we demonstrate that the age-related trends for the harmonized fluid and crystallized composite scores replicate those observed in prior cognitive aging research.

#### 2.3.2 Structural MRI Data

The FreeSurfer 6.0 image analysis suite (http://surfer.nmr.mgh.harvard.edu/; Fischl & Dale, 2000) was used for cortical reconstruction and volumetric segmentation of all the structural MPRAGE images for all four studies. Estimated total intracranial volume was also extracted from the structural images derived from the same automated procedures in FreeSurfer (Buckner et al., 2004). WM hypointensities were labeled automatically, based on FreeSurfer 6.0’s probabilistic procedure (Fischl et al., 2002). Prior to any analyses, WMSA volume was adjusted for estimated total intracranial volume (Buckner et al., 2004) and log-transformed due to its positive skewness.

To alleviate potential concerns regarding our utilization of T1-weighted WM-hypointensity volumes as indices of overall cerebral white matter lesion burden (WMSAs), as opposed to the “gold standard” T2-FLAIR WM-hyperintensity volumes, for a subset of participants who underwent both procedures, we correlated the two volumetric measures. T2-FLAIR lesion-mapping was performed using an unsupervised, single-image method described in (Wetter et al., 2016). An updated, Brain Imaging Data Structure (BIDS)-compatible version of the open-source lesion mapping tool was used (https://github.com/mrfil/lesion-mapper-bids/). Developments were also made to reduce the false positive identification of lesions, by leveraging anatomical segmentations from FreeSurfer, to improve the exclusion of hyperintense voxels from non-white matter regions. This updated software can also separate periventricular from deep white matter lesions, by building masks of contiguous hyperintense voxels radiating out from the ventricles. Additional details on the pulse sequence parameters and lesion mapping procedure can be found in the **supplementary material**.

As shown in the **supplementary material**, the two measures were strongly correlated (*r* = .70), which is similar to the correlation reported by Wei and colleagues (2019) in which they also demonstrated that the two indices of white matter atrophy were highly correlated (*r* = .81) among 56 nondemented elderly individuals. They also demonstrated that both white matter lesion indices exhibited roughly equivalent correlations with other variables of interest, including age and CSF biomarkers of Alzheimer’s pathology. This article and our supplementary analyses strongly support the use of T1-weighted white matter hypointensity volumes as a legitimate index of white matter lesions or WMSAs, in the absence of T2-FLAIR imaging.

#### 2.3.3 Optical Measures of Cerebral Arterial Elasticity

Cerebral arterial elasticity was quantified using an estimate of the shape of the cerebral pulse waveform derived from pulse-DOT (Chiarelli et al., 2017; Fabiani et al., 2014). As reviewed in the introduction, the pulse relaxation function, or PReFx, describes the temporal overlap between the forward and backward pressure waves generated during each cardiac cycle. A greater overlap between these two waves is associated with low PReFx values and indexes arteriosclerosis, whereas a small overlap is associated with high PReFx values and higher arterial elasticity.

To derive PReFx, optical AC intensity data (i.e., the average measures of the amount of light produced by a specific source and reaching a specific detector during a multiplexed 1.6-ms interval) at 830 nm were normalized, movement corrected (Chiarelli et al., 2015), and band-pass filtered between 0.5 and 10 Hz by using a Butterworth filter. The arterial pulse waveform for each channel was obtained by averaging the AC light intensity time-locked to the peak of the R wave of the EKG, ensuring that the same pulse cycle was measured at all locations. Only source-detector distances between 2 and 6 cm were used in the analysis, allowing us to image phenomena occurring approximately between 1 and 3 cm in depth, thereby providing coverage of most of the cerebral cortex. Here, we employed a global measure of cerebral arterial elasticity, which is the average PReFx across all voxels.

#### 2.3.4 Statistical Analyses

##### 2.3.4.1 Piecewise Regression (All Participants)

We conducted four piecewise linear regression analyses in which two lines joined by a single breakpoint were fit to the data, allowing us to estimate the ‘best’ age at which the pre- and post-breakpoint slopes significantly differ. This was done using the *segmented* package (Muggeo, 2008) in R (R Core Team, 2022) and RStudio (RStudio Team, 2020). As a reminder, these analyses were used to estimate a single breakpoint and to test the significance of the difference in slopes between the two fitted lines (i.e., whether the breakpoint did exist) for the relationships between age and PReFx and WMSAs. Identical analyses were conducted with pulse pressure as the vascular index, as opposed to PReFx (**supplementary materials**). High similarity between the breakpoint estimates for age vs. PReFx and age vs. pulse pressure would provide converging evidence from two independent indices of vascular health, thereby increasing confidence in the estimated age range during which vascular problems may accelerate. With respect to the cascade depicted in Figure 1, a large difference in the breakpoint estimates for age vs. PReFx/pulse pressure and age vs. WMSAs, with non-overlapping confidence intervals, may indicate that, on average, precipitous declines in arterial elasticity (arteriosclerosis) *temporally precede* an acceleration in white matter lesion burden, suggesting that these two phenomena may be causally related. Piecewise regression analyses were also conducted between age and estimates of fluid cognition and verbal cognition.

To briefly summarize this analytic approach, via an iterative process, an optimal breakpoint is estimated that minimizes the difference in slope between the two linear regression lines (see Muggeo, 2008 for a detailed explanation of the model and estimation process). We employed the (pseudo) Score statistic (Muggeo, 2016) via the ‘pscore.test()’ function to test for a non-zero difference-in-slope parameter of the bilinear relationship. Simulation studies have shown that this test is more statistically powerful than the alternative Davies test (Davies, 1987) when the existence of only one breakpoint is being tested (Muggeo, 2016), as is done in the present paper. The p-values for the four breakpoint estimates detailed in the main text were adjusted for multiple comparisons using the Holm-Bonferroni procedure.

##### 2.3.4.2 Piecewise Regression (By Sex)

Given extensive evidence that the progression of arteriosclerosis throughout the lifespan generally differs by biological sex, we repeated the previously described piecewise regressions separately for each sex. This equates to eight age-related breakpoint estimates for PReFx, WMSA volume, fluid cognition, and crystallized cognition. These analyses were also repeated with systemic pulse pressure (**supplementary materials)**. The *p*-values for the eight estimates were adjusted for multiple testing using the Holm-Bonferroni procedure.

##### 2.3.4.3 Multiple-Mediator Path Analyses

As described in the introduction, we formed *a priori* hypotheses regarding the relationships among measures representing links in the hypothetical causal chain (**Figure 1**). Specifically, we aimed to test a serial multiple mediator model with two sequential mediators: *X* = age, *M*_1_ = PReFx or Pulse Pressure (Vascular Index), *M*_2_ = WMSA Volume (Structural Index), *Y* = cognitive ability, fluid or crystallized). Consequently, we performed several multiple-mediator mediation analyses with two intervening variables. Using the structural equation modeling package *lavaan* (Rosseel, 2012) in R, we conducted four *two-mediator* path analyses. Missing data was handled via listwise deletion, such that if a given subject has a missing value for at least one measure in a model, it is excluded from all tests of that model. Sex was controlled for as a covariate in all regression-based mediation analyses. Ultimately, we tested the following relationships:

1. Age → PReFx → WMSA Volume → Fluid Cognition *(Sex Controlled as Covariate)*
2. Age → PReFx → WMSA Volume → Crystallized Cognition *(Sex Controlled as Covariate)*
3. Age → Pulse Pressure → WMSA Volume → Fluid Cognition *(Sex Controlled as Covariate)*
4. Age → Pulse Pressure → WMSA Volume → Crystallized Cognition *(Sex Controlled as Covariate)*

Note that we did not predict a significant relation between PReFx or pulse pressure and crystallized ability or a significant indirect effect with WMSAs as the mediator of this relationship, because crystallized abilities are largely invulnerable to normal neurocognitive decline (until ∼70 years of age) and continue to increase throughout adulthood. Therefore, this model was included to demonstrate a dissociation between the effects of age-related changes in our two intervening variables on fluid abilities versus crystallized abilities. Models 3 and 4, in which pulse pressure serves as the vascular index, are presented in the **supplementary materials**.

The cascade model depicted in Figure 1 leads to the *a priori* hypotheses that an age-related loss of cerebral arterial elasticity may *indirectly* affect cognition via its deleterious impact on white matter integrity, manifesting in part as demyelination and axonal loss.

Because the distribution of an indirect effect (the product of the relevant path coefficients; *a_1_ × b_1_*, *a_2_ × b_2_*, or *a_1_ × d_21_ × b_2_*) is typically non-normal and asymmetric (Hayes, 2009), a bootstrap approach was implemented for significance testing of the indirect effects in each model, based on 5,000 bootstrap samples (Preacher & Kelley, 2011). To clarify, in a two-mediator model, path *d_21_* corresponds to the effect of *M1* on *M2* when controlling for *X.* We also report coefficient estimates for the total effect, *c* (*Y* regressed on *X* without covariates), and direct effect, *c’*, which is the effect of *X* on *Y* when statistically controlling for the effect of both mediators, *M_1_* and *M_2_* on *Y* (paths *b_1_* and *b_2_*). As the primary outcome of interest for each mediation analysis was the two-mediator sequential indirect effect, the bootstrapped p-values for each two-mediator indirect effect were adjusted using the Holm-Bonferroni approach, once again.

## 3. RESULTS

### 3.1 Piecewise Regression Analyses (Overall)

As predicted, the relationships between age and arterial health, and age and white matter lesions exhibited non-linear relationships, which were generally well-described by bilinear trends (**Figure 2**).

**Figure 2:**
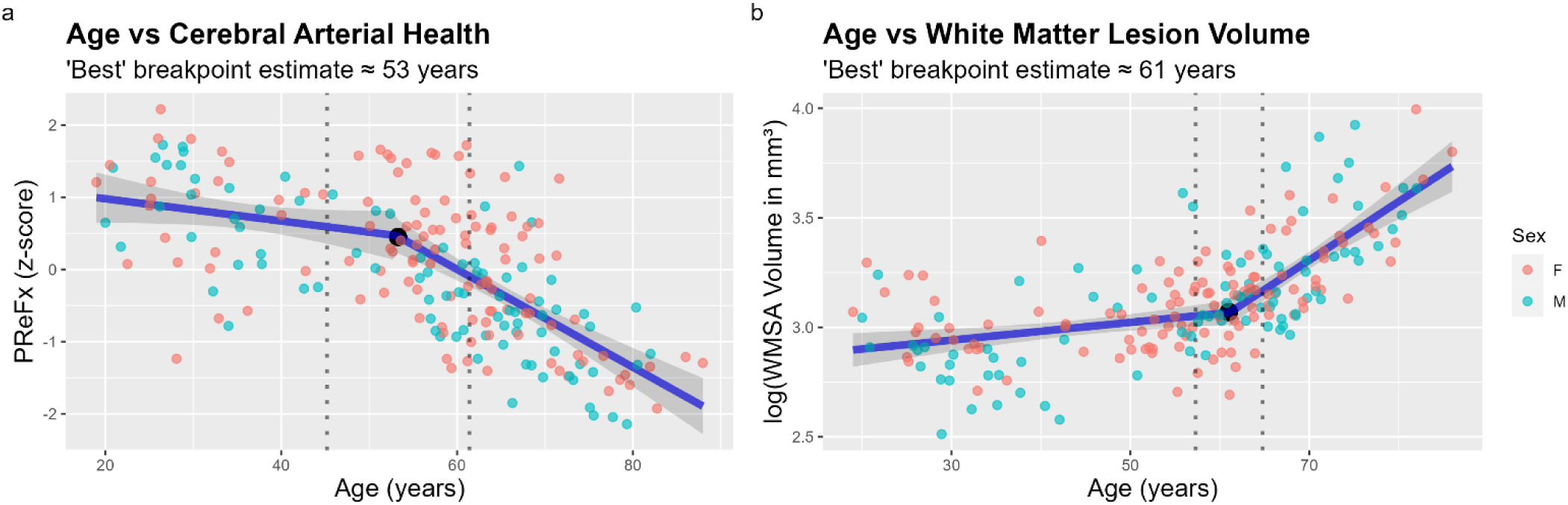
Scatterplots with piecewise linear regression lines superimposed (shaded areas indicate 95% confidence interval bands). Participants are color-coded by their self-reported sex (females = peach, males = aqua). The black dot in each plot indicates the ‘best’ breakpoint estimate for each piecewise relationship, or the age at which the difference in slopes between the two regression lines is largest—these breakpoints are statistically significant after employing a Holm-Bonferroni correction. The vertical dashed lines represent the lower and upper limits of the 95% confidence intervals for each estimate. The breakpoint estimates for **(a)** cerebral arterial health and **(b)** cerebral white matter atrophy differ by approximately 8 years, with partially nonoverlapping confidence intervals. Together, this suggests that, on average, declines in cerebrovascular health precede the accelerated formation of white matter lesions. WMSA = white matter signal abnormalities.

#### 3.1.1 Age vs. Pulse Relaxation Function (Cerebral Arterial Elasticity)

Piecewise linear regression analyses of the association between age and PReFx (*n* = 203) yielded a single breakpoint estimate (age = 53.09 years, *SE* = 4.05, *p*_Holm_ < .001; 95% CI [45.11, 61.08]), indicating that the difference in slopes was statistically significant before and after this point (*β*_1_ = -.25, 95% CI [-.54, .04]; *β*_2_ = -1.10, 95% CI [-1.37, -.84]; standardized regression coefficients reported), with only the post-breakpoint slope (*β*_2_) indicating a significant relationship (**Figure 2a**). As such, our data suggest that a reliable, precipitous decline in cerebral arterial elasticity may not occur until late middle age. The results for systemic pulse pressure virtually replicate those of PReFx: (age = 51.29 years, *SE* = 3.58, *p* < .001; 95% CI [44.25, 58.34]).

#### 3.1.2 Age vs. White Matter Signal Abnormality Volume (White Matter Lesion Load)

Piecewise linear regression analyses of age and WMSA volume (*n* = 202) yielded a single breakpoint estimate (age = 61.09 years, *SE* = 1.92, *p*_Holm_ < .001; 95% CI [57.31, 64.87]), indicating that slopes were reliably different before and after this point (*β*_1_ = .07, 95% CI [.02 .11]; *β*_2_ = .44, 95% CI [.33, .54]; standardized regression coefficients reported), with slope *β*_2_ larger than slope *β*_1_ (**Figure 2b**), indicating accelerated white matter atrophy after this age. Note that this breakpoint estimate is larger than that the PReFx and pulse pressure estimates, and its 95% confidence interval only partially overlaps that of the vascular estimates, suggesting that declines in vascular health generally precede the accelerated formation of white matter lesions.

#### 3.1.3 Age vs. Fluid and Crystallized Cognition

Piecewise linear regression analyses of age and fluid cognition (*n* = 194) yielded a single breakpoint estimate (age = 55.26 years, *SE* = 5.19, *p*_Holm_ < .01; 95% CI [45.02, 65.05]), indicating that slopes were reliably different before and after this point (*β*_1_ = -.29, 95% CI [-.57 -.02]; *β*_2_ = -.96, 95% CI [-1.26, -.65]; standardized regression coefficients reported), with slope *β*_2_ larger than slope *β*_1_ (**Figure 3a**), indicating average accelerated declines in fluid abilities such as processing speed, working memory, and reasoning with age.

**Figure 3:**
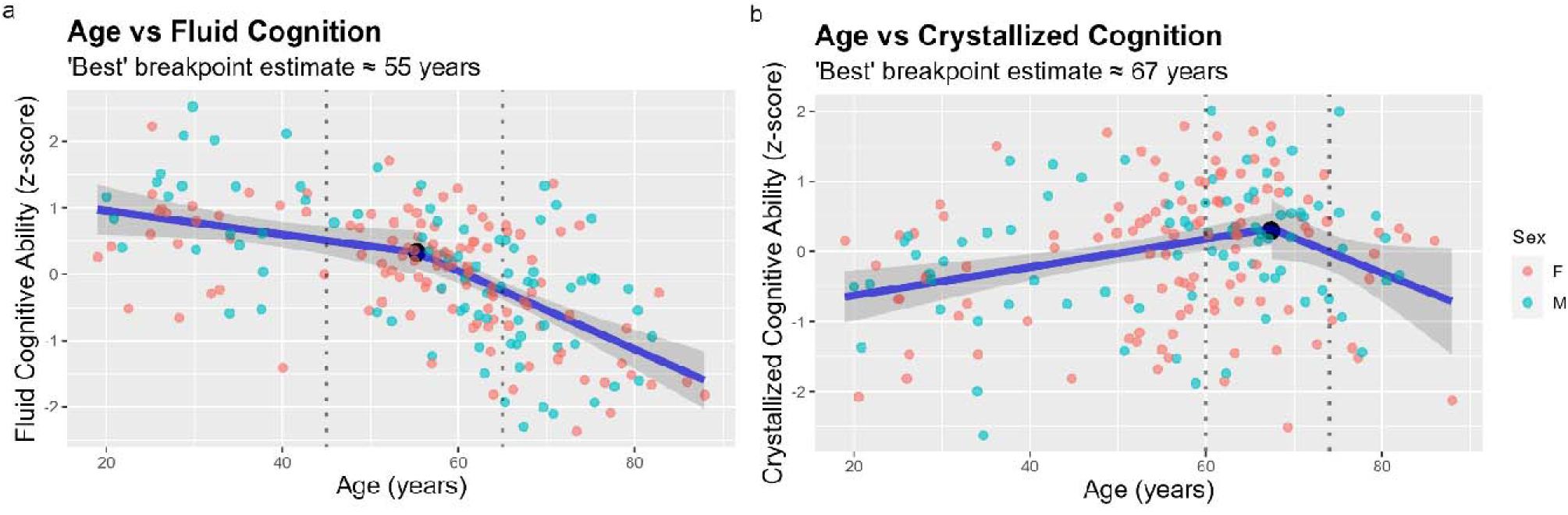
Scatterplots with piecewise linear regression lines superimposed (shaded areas indicated 95% confidence interval bands). Participants are color-coded by their self-reported sex (females = peach, males = aqua). The black dot in each plot indicates the ‘best’ breakpoint estimate for each piecewise relationship, or the age at which the difference in slopes between the two regression lines is largest—these breakpoints are statistically significant after employing a Holm-Bonferroni correction. The vertical dashed lines represent the lower and upper limits of the 95% confidence intervals for each estimate. The breakpoint estimates for **(a)** fluid cognition and **(b)** crystallized or verbal cognition differ are seemingly attributable to different physiological phenomena, due to their relatively large discrepancy. The accelerated decline in fluid cognition seemingly occurs within the interval during which we observe accelerated declines in cerebrovascular health and white matter health in our data, which is consistent with prior evidence indicating that fluid abilities are more susceptible to neurobiological deterioration than crystallized cognition.

Piecewise linear regression analyses of age and crystallized cognition (*n* = 194) yielded a single breakpoint estimate (age = 67.40 years, *SE* = 3.52, *p*_Holm_ < .01; 95% CI [60.46, 74.34]), indicating that slopes were reliably different before and after this point (*β*_1_ = .33, 95% CI [.15 .50]; *β*_2_ = -.82, 95% CI [-1.66, .02]; standardized regression coefficients reported), with only slope *β*_1_ statistically significant (**Figure 3b**). The absence of a significant post-breakpoint slope is likely attributable to the small number of participants older than 67 years of age and resultant lack of statistical power for this portion of the analyses.

### 3.2 Piecewise Regression Analyses (By Sex)

The following results illustrate sex-related differences in the trajectories of various neurophysiological and cognitive outcomes, as demonstrated in **Figures 4 and 5**.

**Figure 4:**
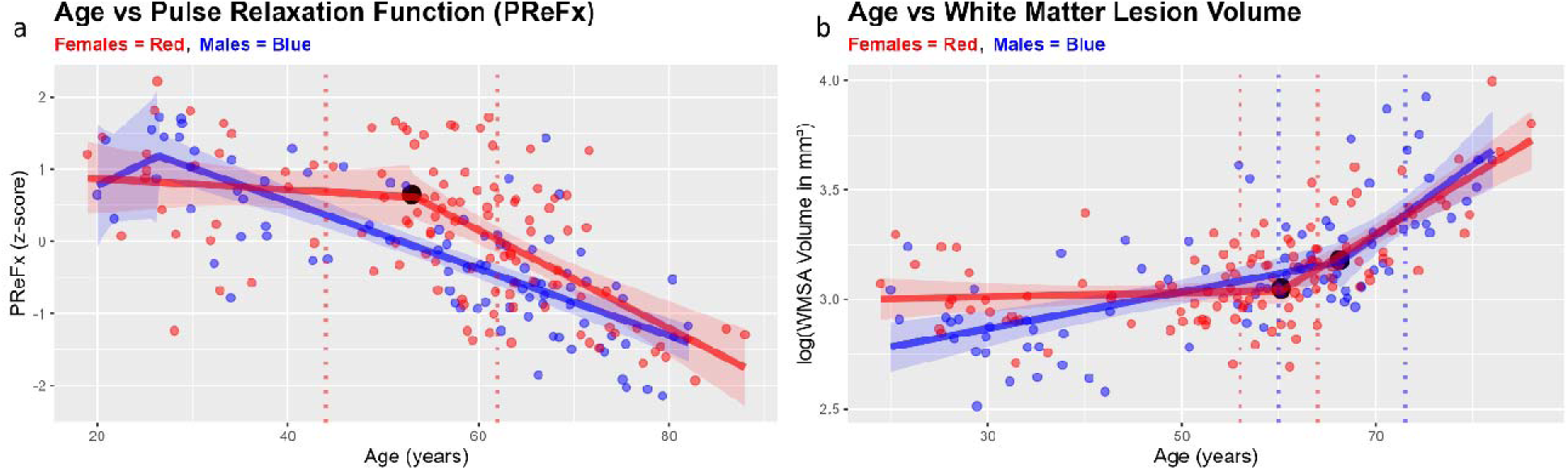
Scatterplots with separate piecewise linear regression lines for females and males superimposed (red and blue shaded areas indicated 95% confidence interval bands for females and males, respectively). The black dot(s) in each plot indicates the ‘best’ breakpoint estimate for each piecewise relationship, or the age at which the difference in slopes between the two regression lines is largest—these breakpoints are statistically significant after employing a Holm-Bonferroni correction. The red and blue vertical dashed lines represent the lower and upper limits of the 95% confidence intervals for each estimate for females and males, respectively. The breakpoint estimates for **(a)** PReFx are only significant for females at age 53, roughly coinciding with the onset of menopause and a drop in estrogen production. Males’ cerebrovascular health seemingly declines steadily throughout the lifespan. Males and females both exhibited statistically significant breakpoints for **(b)** white matter lesion volume, with females’ occurring at 60 and males’ occurring later at age 66. Contrary to males, however, females’ white matter health does not significantly decrease prior to age 60, presumably due to the loss of the vasoprotective effects of estradiol after age 50, which then manifest as white matter lesions years later. These results are consistent with the hypothesis that cerebrovascular declines contribute to the formation of white matter lesions.

**Figure 5:**
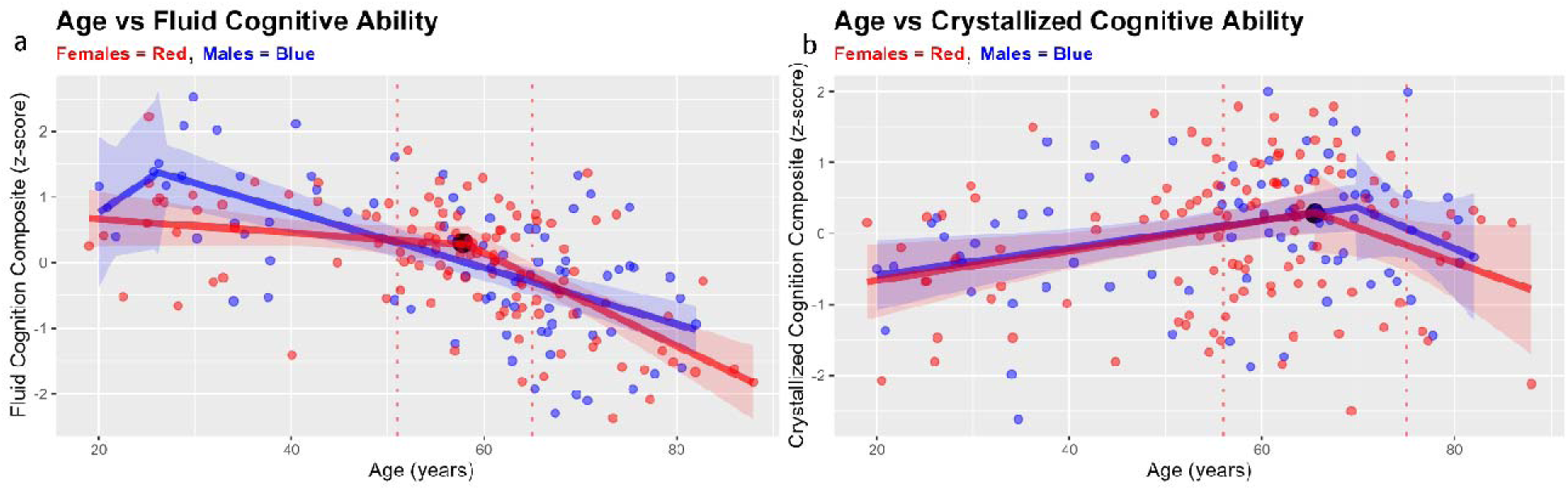
Scatterplots with separate piecewise linear regression lines for females and males superimposed (red and blue shaded areas indicated 95% confidence interval bands for females and males, respectively). The black dot(s) in each plot indicates the ‘best’ breakpoint estimate for each piecewise relationship, or the age at which the difference in slopes between the two regression lines is largest—these breakpoints are statistically significant after employing a Holm-Bonferroni correction. The red and blue vertical dashed lines represent the lower and upper limits of the 95% confidence intervals for each estimate for females and males, respectively. The breakpoint estimates for **(a)** fluid cognition are only significant for females at age 58, which is within the presumed interval during which vascular health declines and white matter lesion development accelerates. Males’ fluid abilities seemingly decline steadily throughout the lifespan. Only females both exhibited statistically significant breakpoints for **(b)** crystallized or verbal cognition, with females’ occurring at 65. This suggests that crystallized cognition may be less vulnerable to declines in cerebrovascular health and cerebral matter atrophy.

#### 3.2.1 Sex-Related Differences in Cerebral Arteriosclerosis (PReFx)

Piecewise linear regression analyses of the association between age and PReFx (**Figure 4a**) yielded a significant breakpoint estimate for females (age = 53.29 years, *SE* = 4.54; 95% CI [44.29, 62.29], *p*_Holm_ < .01) but not for males (age = 26.44 years, *SE* = 4.60; 95% CI [17.29, 35.59], *p*_Holm_ = .27), suggesting that women’s cerebrovascular health, on average, is minimally affected until late middle-age whereas men’s declines steadily, possibly beginning during young adulthood.

#### 3.2.2 Sex-Related Differences in White Matter Lesion Formation (WMSA Volume)

Piecewise linear regression analyses of the association between age and WMSA volume (**Figure 4b**) yielded significant breakpoint estimates for females (age = 60.21 years, *SE* = 1.88; 95% CI [56.48, 63.93], *p*_Holm_ < .001) and males (age = 66.26 years, *SE* = 3.25; 95% CI [59.80, 72.72], *p*_Holm_ < .01), which were separated by partially non-overlapping 95% confidence intervals. It’s also important to note that, for males, both the pre- and post-breakpoint slopes are significant, which is consistent with the steady decline in cerebrovascular health that begins during young adulthood, as shown in **Figure 4a**. However, for women, only the post-breakpoint slope is statistically significant, which mirrors women’s age-related vascular trajectory, as the pre-breakpoint slope is effectively zero for PReFx and WMSA volume. Moreover, this provides support for the hypothesis that females’ vascular health is largely protected until middle age (i.e., the onset of menopause), likely due to the anti-inflammatory and vasoprotective effects of estrogens.

#### 3.2.3 Sex-Related Differences in Cognitive Decline (Fluid and Crystallized Ability)

Piecewise linear regression analyses of the association between age and fluid cognition (**Figure 5a**) yielded a significant breakpoint estimate for females (age = 58.28 years, *SE* = 3.50; 95% CI [51.34, 65.22], *p*_Holm_ < .001) but not for males (age = 26.20 years, *SE* = 4.81; 95% CI [16.63, 35.77], *p*_Holm_ = .69). Once again, the presumed temporal onset of precipitous declines in fluid cognition is highly similar to the estimated onset of accelerated white matter lesion formation (60 years), which is also strongly negatively correlated with fluid cognition (*r*(191) = -.48, *p* < .001). Also, males exhibit a relatively steady decline in fluid cognition, which is commensurate with the relatively linear trends observed for vascular and white matter degradation in men. These results are consistent with a model that posits declines in cerebrovascular health contribute to declines in fluid abilities during aging.

Piecewise linear regression analyses of the association between age and crystallized cognition (**Figure 5b**) yielded a significant breakpoint estimate for females (age = 65.50 years, *SE* = 4.77; 95% CI [56.0, 74.95], *p*_Holm_ < .05) but not for males (age = 69.77 years, *SE* = 5.60; 95% CI [58.62, 80.92], *p*_Holm_ = .27). The lag associated with precipitous declines in crystallized cognition (compared to fluid cognition) likely reflects the contributions of a semi-independent pathophysiological mechanism, including declines in medial temporal lobe/hippocampal volume and formation of Alzheimer’s disease-related pathologies such as β-amyloid and tau deposition (McDonough et al., 2016; Villeneuve & Jagust, 2015), as opposed to the formation of white matter lesions.

### 3.3 Multiple-Mediator Path Analyses

#### 3.3.1 Age → PReFx → WMSA Volume → Cognition (Fluid and Crystallized)

To complement the prior analyses, we conducted several multiple-mediator path analyses to simultaneously test the hypothesized relationships among age, arterial health (PReFx or systemic pulse pressure), white matter lesions (WMSA volume), and cognition (fluid or crystallized). As a reminder, sex was included as a covariate in these analyses, but is not shown in the mediation diagrams to minimize visual clutter. Firstly, we tested the two-mediator indirect effect of age on fluid cognition via PReFx and WMSA volume (**Figure 6**; *n* = 193). Based on a bias-corrected bootstrapped 95% confidence interval, the two-mediator serial indirect effect (*a_1_ × d_21_ × b_2_*) was significant, suggesting that age-related declines in cerebral arterial health impact fluid abilities *via* their deleterious effects on white matter integrity. Importantly, we observed a significant indirect effect of age on fluid cognition *via* white matter lesions (*a_2_× b_2_*), but *not via* cerebral arterial health (*a_1_ × b_1_*), suggesting, once again, that age-related declines in cerebral health largely manifest as degraded white matter, which negatively impacts fluid abilities such as processing speed, working memory, and reasoning. Lastly, no significant indirect effects were observed when crystallized cognition was the final outcome (**Figure 7**). The same pattern of significant results was observed when pulse pressure was used as the vascular index, instead of PReFx, supporting our findings (see **supplementary material**).

**Figure 6:**
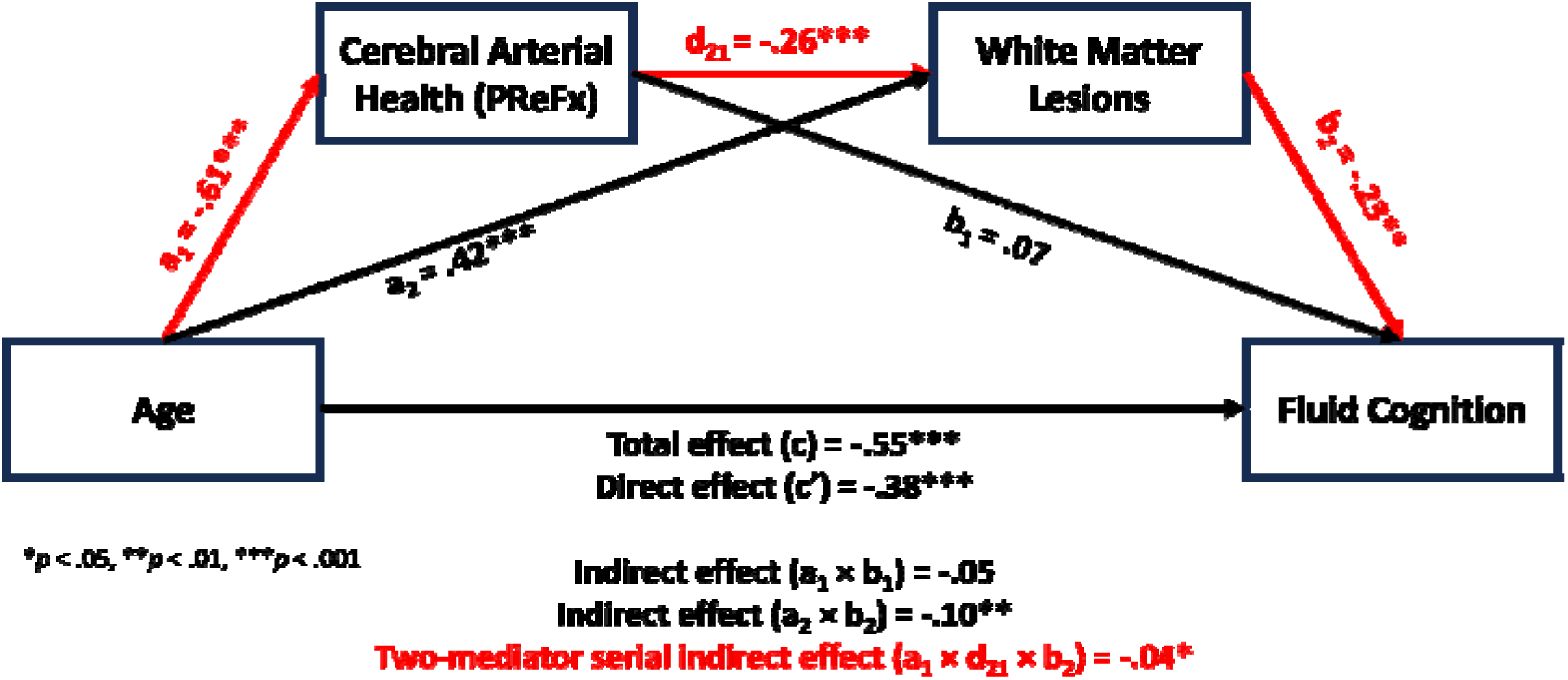
Serial mediation diagram examining the effect of age on fluid cognition via two intervening variables, PReFx (a pulse-DOT parameter measuring cerebral arterial stiffness) and T1w white matter hypointensities (an index of white matter lesions or WMSAs). Self-reported sex is controlled for in these analyses but is not included in the diagram to maintain visual clarity. The two-mediator sequential indirect effect is significant, which is consistent with the hypothesis that age-related declines in cerebral arterial health contribute to the formation of white matter lesions and negatively impact fluid cognition. Red font/arrows indicate paths implicated in the multiple-mediator indirect effect. \**p* < .05, \*\**p* < .01, \*\*\**p* < .001

**Figure 7:**
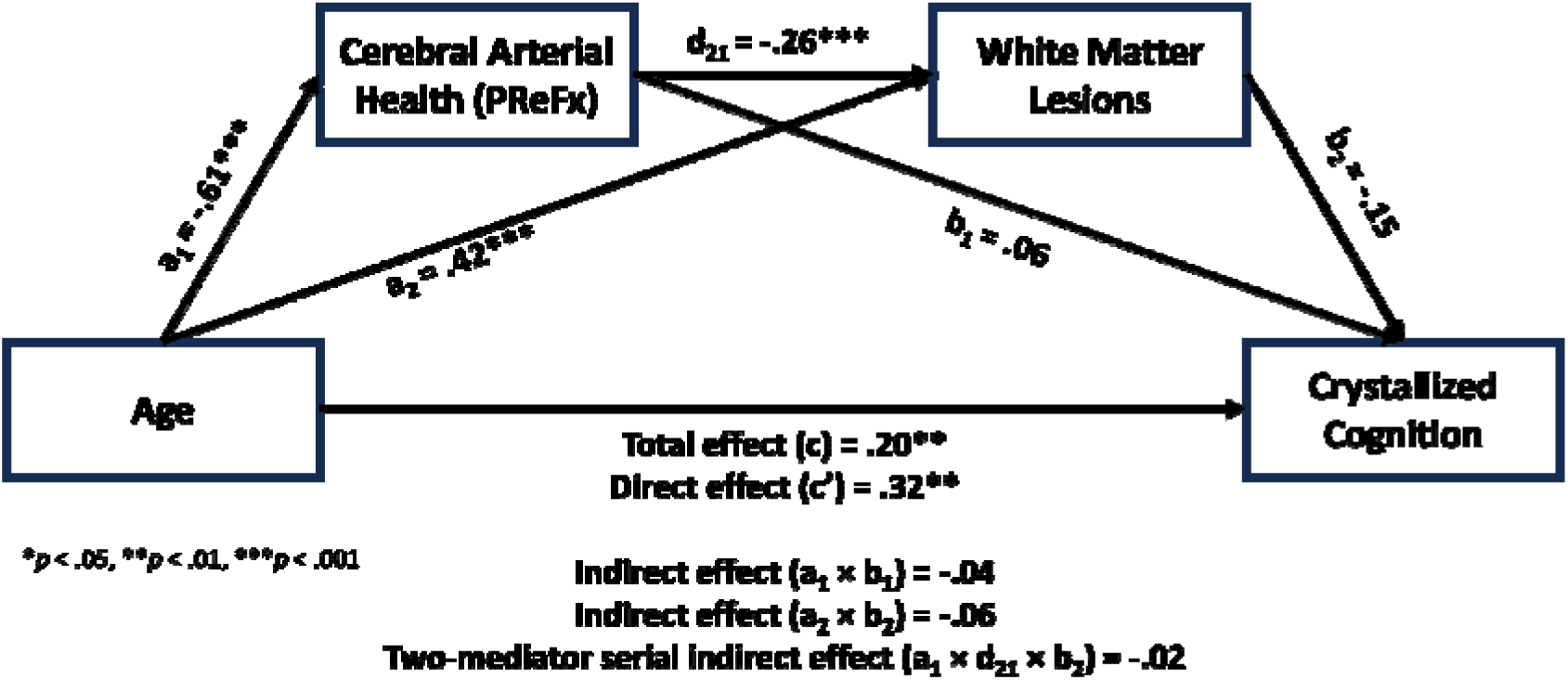
Serial mediation diagram examining the effect of age on crystallized cognition via two intervening variables, PReFx (a pulse-DOT parameter measuring cerebral arterial stiffness) and T1w white matter hypointensities (an index of white matter lesions or WMSAs). Self-reported sex is controlled for in these analyses but is not included in the diagram to maintain visual clarity. The two-mediator sequential indirect effect is not significant, which is consistent with the hypothesis that age-related declines in cerebral arterial health contribute to the formation of white matter lesions but do not severely impact crystallized cognition. \**p* < .05, \*\**p* < .01, \*\*\**p* < .001

## 4. DISCUSSION

### 4.1 General Discussion

We have employed a novel approach to test a cascade model of the role of arteriosclerosis in cognitive aging and ultimately dementia (**Figure 1**). This approach includes the use of piecewise correlations to establish the relative timing of emergence of phenomena in the cascade at the group level and thus enable Granger-like causal inferences in an otherwise cross-sectional sample. Based on two largely independent measures of cerebral (pulse-DOT) and systemic (pulse pressure) arterial health, we have provided converging evidence that group-level age-related declines in arterial elasticity appear to precede accelerated white matter atrophy by 5-10 years and therefore potentially contribute to the formation of white matter lesions, which, in turn, negatively impact fluid, but not crystallized abilities. Specifically, we observed a significant two-mediator indirect effect of age on fluid cognition via cerebral arterial health (measured via PReFx) and white matter lesions such that age-related declines in cerebral arterial health lead to increases in white matter lesions that negatively impact performance on tests of fluid abilities, which is consistent with the temporal ordering of mechanisms posited in the hierarchical model (see **Figure 1**). Also, as expected, crystallized abilities remained relatively intact despite these age-related neurovascular declines, which is consistent with evidence showing that such abilities do not decline until even older age (∼70 years). Similarly, piecewise linear regression analyses yielded breakpoint estimates of age that were separated by approximately eight years for cerebral arterial health (∼53 years) and WMSAs (∼61 years), although the 95% confidence intervals for the two estimates partially overlapped. As reported in the **supplementary materials**, we replicated the results of all analyses using an independent and readily accessible estimate of arterial health, systemic pulse pressure, thereby providing converging evidence for the mediating role of cerebrovascular mechanisms in cognitive aging.

Of extreme importance, we also observed significant sex-related differences in the age-related progression of cerebral arteriosclerosis, white matter degradation, and declines in fluid cognition, all of which are consistent with the proposed sequence and mediating role of cerebrovascular health (see **Figure 1**). As women’s estrogen production drops during and after menopause (around age 50), this ultimately mitigates the vascular protective effects of estrogens such as estradiol, thereby accelerating arterial stiffness and white matter lesion formation, which may deleteriously impact fluid cognition, but not crystallized cognition (or at least, to a lesser extent).

### 4.2 Sex-Related Differences in Age-Related Neurovascular Decline

Overall, our results suggest that age-related declines in indices of vascular integrity occur earlier in males than females, as indicated by piecewise regression analyses, possibly due to the vascular protective effects of estrogen, which are largely lost after menopause and related decline in estrogen production (Hurn et al., 1995). This sex-related difference in the progression of arteriosclerosis *may* affect the onset of acceleration in white matter lesion development in males versus females, as evidenced by a later inflection point estimate for males (**Figure 4b**), although a significant steady decline in cerebral white matter health begins in young adulthood for men, but not women.

Prior cross-sectional research has suggested that the average pulse pressure of women is lower than that of men earlier in life (which we also observed; see **supplementary materials**), but typically exhibits an accelerated increase after age 50 that is driven by increases in systolic blood pressure (Chou et al., 2022; Skurnick et al., 2010). This is consistent with our results, as we observed a significant, precipitous average increase in pulse pressure for women during age 50 (around the onset of menopause; National Institute on Aging, 2021) whereby women’s mean pulse pressure eventually exceeds men’s mean pulse pressure (cross-over effect; **Figure S4**). However, longitudinal data indicate that the mean pulse pressure of middle-aged to older women exhibits a similar slope to or converges with men’s mean pulse pressure, but does not exceed it, resulting in the absence of a sex-related cross-over effect (Franklin et al., 1999; Pearson et al., 1997). Researchers have hypothesized that this discrepancy in cross-sectional versus longitudinal findings may be due to selective mortality in men with elevated pulse pressure.

Piecewise analyses of PReFx (cerebral arterial elasticity) and age demonstrated similar trends to pulse pressure such that a precipitous decline in PReFx occurs later in females than males. For PReFx, the inflection point for females, which was significant after correction for multiple comparisons, was estimated to occur around age 53, while males did not exhibit a significant inflection point, with a steep decrease in PReFx beginning in young adulthood. Using transcranial Doppler ultrasonography and a breath holding manipulation to assess cerebrovascular reactivity in middle-aged men and women, Matteis et al. (1998) observed significantly lower breath-holding indices (indicating *lower* cerebral vasoreactivity) in postmenopausal women than premenopausal women, younger men, and older men. Breath-holding indices were also significantly lower in postmenopausal than premenopausal women of the same age, all of which strongly suggest that the loss of estradiol during menopause deleteriously affects women’s cerebrovascular health. Additional evidence suggests that administration of estradiol in postmenopausal women partly mitigates the clinical symptomatology of coronary heart disease (Reis et al., 1994). Moreover, an increase in cerebral arterial pulsatility in postmenopausal women is significantly correlated with menopausal age (i.e., age since menopausal onset) but not chronological age, with postmenopausal women displaying higher pulsatility values than premenopausal women of the same age (Penotti et al., 1996), supporting the claim that estrogen confers protective benefits to the cerebral vasculature.

The relationship between age and white matter lesion volume, based on piecewise regression analyses, again suggested that a significant decline in white matter integrity occurs later in females than in males. Importantly, the inflection point estimate for females (∼60 years of age) temporally succeeds the estimates of the inflection points for the vascular measures by 7-8 years, which is consistent with the claim that declines in vascular health partly contribute to declines in the brain’s structural integrity, particularly when manifested as white matter lesions (Pantoni & Garcia, 1997).

### 4.3 Limitations and Future Directions

Although the analytic approach used in this study has the advantage of allowing some temporal Granger-like inferences in an otherwise cross-sectional sample, it also has a number of limitations. Firstly, because regression-based mediation analyses assume linear relationships between predictors and outcomes, when including age and PReFx/pulse pressure or WMSAs in a mediation model, statistical power is likely reduced, as these relationships are not best described by a linear function. Therefore, it is arguable that the statistical power of our mediation analyses suffers, suggesting that the mediating roles of arteriosclerosis and white matter lesion formation are actually more robust than that reported here. Secondly, due to possible cohort effects that are not unique to this study, the utilization of a cross-sectional design, as opposed to a longitudinal design, limits the strength of causal inferences regarding putative mechanisms.

To bolster the current findings, future studies should employ even larger sample sizes and, ideally, longitudinal designs, thereby permitting stronger causal inferences regarding neurocognitive aging mechanisms. Also, other significant outcomes, such as BOLD signal-based measures of cerebrovascular reactivity that capture the integrity of neurovascular coupling (Tarantini et al., 2017), may serve as complementary indices of cerebrovascular health when formally testing the hierarchical model. Moreover, the inclusion of latent variables via structural equation modeling, estimated based on multiple reliable indices (i.e., observed variables) representing each link or latent construct in the casual chain, may provide a more comprehensive and accurate characterization of neurocognitive aging. Additionally, it will be important to examine the regional, as opposed to global, cerebral variations in arterial health afforded by PReFX. It will also be important to examine the mitigating effects of cardiorespiratory fitness as a way of mitigating and/or slowing the negative effects of age-related declines in cerebrovascular health.

### 4.3 Conclusions

To conclude, an expanded sample allowed us to investigate the putative mechanisms of neurocognitive aging (**Figure 1**) that often result in mild cognitive impairment or even dementia, dramatically impairing a person’s quality of life and ability to function in the world. We elaborated upon preliminary correlational analyses presented by Kong et al. (2020) by ***a)*** increasing the sample size more than four-fold, ***b)*** employing piecewise regression analyses to estimate ages or age ranges during which arteriosclerosis (cerebral arterial stiffening) and white matter atrophy accelerate, ***c)*** conducting several multiple-mediator path analyses to further substantiate the hypothesized causal links posited in the hierarchical model of neurocognitive aging, ***d)*** identifying clinically relevant and robust sex-related differences in aging trajectories of cerebrovascular health, white matter integrity, and fluid cognition, and ***d)*** replicating all analyses involving PReFx (which provides a direct measure of cerebrovascular health) using an independent, peripheral index of vascular health, systemic pulse pressure (see **supplementary materials** for these confirmatory results). Findings were also consistent with the proposed sequence of events leading to cognitive decline such that average age-related declines in vascular health preceded the acceleration in appearance of white matter lesions. In addition, accelerated declines in fluid cognition roughly coincided with accelerated declines in white matter health, while crystallized cognition was spared until nearly age 70. Similarly, results of multiple-mediator path analyses (controlling for sex) were consistent with the proposed sequence of events in neurocognitive aging.

## Acknowledgments

This work is part of the dissertation of the first author, in partial fulfillment of the PhD requirements. The research was supported by NIA grants R01AG059878 and RF1AG062666 to M. Fabiani and G. Gratton and by NICCH grant 5R90AT008924 to B. Zimmerman. We acknowledge the support of the Bioimaging Center of the Beckman Institute for Advanced Science and Technology at the University of Illinois Urbana-Champaign (UIUC-BI-BIC).

## REFERENCES

Aging, N. I. on. (2021). What is MenoPause? https://www.nia.nih.gov/health/what-menopause

Beck, A. T., Steer, R. A., & Brown, G. K. (1996). Manual for the Beck Depression Inventory-II.

Buckner, R. L., Head, D., Parker, J., Fotenos, A. F., Marcus, D., Morris, J. C., & Snyder, A. Z. (2004). A unified approach for morphometric and functional data analysis in young, old, and demented adults using automated atlas-based head size normalization: reliability and validation against manual measurement of total intracranial volume. NeuroImage, 23(2), 724–738. 10.1016/j.neuroimage.2004.06.018

Burt, V. L., & Harris, T. (1994). The third National Health and Nutrition Examination Survey: contributing data on aging and health. The Gerontologist, 34(4), 486–490. 10.1093/geront/34.4.486

Calamante, F., Thomas, D. L., Pell, G. S., Wiersma, J., & Turner, R. (1999). Measuring Cerebral Blood Flow Using Magnetic Resonance Imaging Techniques. Journal of Cerebral Blood Flow & Metabolism, 19(7), 701–735. 10.1097/00004647-199907000-00001

Chiarelli, A. M., Fletcher, M. A., Tan, C. H., Low, K. A., Maclin, E. L., Zimmerman, B., Kong, T., Gorsuch, A., Gratton, G., & Fabiani, M. (2017). Individual differences in regional cortical volumes across the life span are associated with regional optical measures of arterial elasticity. NeuroImage, 162, 199–213. 10.1016/j.neuroimage.2017.08.064

Chiarelli, A. M., Maclin, E. L., Fabiani, M., & Gratton, G. (2015). A kurtosis-based wavelet algorithm for motion artifact correction of fNIRS data. NeuroImage, 112, 128–137. 10.1016/j.neuroimage.2015.02.057

Chojdak-Łukasiewicz, J., Dziadkowiak, E., Zimny, A., & Paradowski, B. (2021). Cerebral small vessel disease: A review. Advances in Clinical and Experimental Medicine : Official Organ Wroclaw Medical University, 30(3), 349–356. 10.17219/acem/131216

Chou, C.-H., Yin, J.-H., Lin, Y.-K., Yang, F.-C., Chu, T.-W., Chuang, Y. C., Lin, C. W., Peng, G.-S., & Sung, Y.-F. (2022). The optimal pulse pressures for healthy adults with different ages and sexes correlate with cardiovascular health metrics. Frontiers in Cardiovascular Medicine, 9, 930443. 10.3389/fcvm.2022.930443

Colcombe, S., & Kramer, A. F. (2003). Fitness effects on the cognitive function of older adults: a meta-analytic study. Psychological Science, 14(2), 125–130. 10.1111/1467-9280.t01-1-01430

Corrigan, J. D., & Hinkeldey, N. S. (1987). Relationships between Parts A and B of the Trail Making Test. Journal of Clinical Psychology, 43(4), 402–409. 10.1002/1097-4679(198707)43:4<402::AID-JCLP2270430411>3.0.CO;2-E

Davies, R. B. (1987). Hypothesis testing when a nuisance parameter is present only under the alternative. Biometrika, 74(1), 33–43. 10.1093/biomet/74.1.33

Fabiani, M., Asnakew, B. A., Bowie, D. C., Chism, S. M., Clements, G. M., Gardner, J. C., Islam, S. S., Rubenstein, S. L., & Gratton, G. (2022). A healthy mind in a healthy body: Effects of arteriosclerosis and other risk factors on cognitive aging and dementia. In K. D. Federmeier & B. R. B. T.-P. of L. and M. Payne (Eds.), Psychology of Learning and Motivation (Vol. 77, pp. 69–123). Academic Press. 10.1016/bs.plm.2022.08.001

Fabiani, M., Low, K. A., Tan, C.-H., Zimmerman, B., Fletcher, M. A., Schneider-Garces, N., Maclin, E. L., Chiarelli, A. M., Sutton, B. P., & Gratton, G. (2014). Taking the pulse of aging: mapping pulse pressure and elasticity in cerebral arteries with optical methods. Psychophysiology, 51(11), 1072–1088. 10.1111/psyp.12288

Fischl, B, & Dale, A. M. (2000). Measuring the thickness of the human cerebral cortex from magnetic resonance images. Proceedings of the National Academy of Sciences of the United States of America, 97(20), 11050–11055. 10.1073/pnas.200033797

Fischl, Bruce, Salat, D. H., Busa, E., Albert, M., Dieterich, M., Haselgrove, C., van der Kouwe, A., Killiany, R., Kennedy, D., Klaveness, S., Montillo, A., Makris, N., Rosen, B., & Dale, A. M. (2002). Whole brain segmentation: automated labeling of neuroanatomical structures in the human brain. Neuron, 33(3), 341–355. 10.1016/s0896-6273(02)00569-x

Franklin, S S, Khan, S. A., Wong, N. D., Larson, M. G., & Levy, D. (1999). Is pulse pressure useful in predicting risk for coronary heart Disease? The Framingham heart study. Circulation, 100(4), 354–360. 10.1161/01.cir.100.4.354

Franklin, Stanley S. (2006). Hypertension in Older People: Part 1. The Journal of Clinical Hypertension, 8(6), 444–449. 10.1111/j.1524-6175.2006.05113.x

Gratton, G., Chiarelli, A. M., & Fabiani, M. (2017). From brain to blood vessels and back: a noninvasive optical imaging approach. Neurophotonics, 4(3), 31208. 10.1117/1.NPh.4.3.031208

Hedden, T., & Gabrieli, J. D. E. (2004). Insights into the ageing mind: A view from cognitive neuroscience. Nature Reviews Neuroscience, 5(2), 87–96. 10.1038/nrn1323

Hillman, C. H., Erickson, K. I., & Kramer, A. F. (2008). Be smart, exercise your heart: exercise effects on brain and cognition. Nature Reviews Neuroscience, 9, 58–65.

Hurn, P. D., Littleton-Kearney, M. T., Kirsch, J. R., Dharmarajan, A. M., & Traystman, R. J. (1995). Postischemic cerebral blood flow recovery in the female: effect of 17 beta-estradiol. Journal of Cerebral Blood Flow and Metabolism : Official Journal of the International Society of Cerebral Blood Flow and Metabolism, 15(4), 666–672. 10.1038/jcbfm.1995.82

Izzo, J. L. J., & Shykoff, B. E. (2001). Arterial stiffness: clinical relevance, measurement, and treatment. Reviews in Cardiovascular Medicine, 2(1), 29–34,37-40.

Kong, T. S., Gratton, C., Low, K. A., Tan, C. H., Chiarelli, A. M., Fletcher, M. A., Zimmerman, B., Maclin, E. L., Sutton, B. P., Gratton, G., & Fabiani, M. (2020). Age-related differences in functional brain network segregation are consistent with a cascade of cerebrovascular, structural, and cognitive effects. Network Neuroscience (Cambridge, Mass.), 4(1), 89–114. 10.1162/netn_a_00110

Li, Q., Yang, Y., Reis, C., Tao, T., Li, W., Li, X., & Zhang, J. H. (2018). Cerebral Small Vessel Disease. Cell Transplantation, 27(12), 1711–1722. 10.1177/0963689718795148

Mahmood, S. S., Levy, D., Vasan, R. S., & Wang, T. J. (2014). The Framingham Heart Study and the epidemiology of cardiovascular disease: a historical perspective. Lancet (London, England), 383(9921), 999–1008. 10.1016/S0140-6736(13)61752-3

Matteis, M., Troisi, E., Monaldo, B. C., Caltagirone, C., & Silvestrini, M. (1998). Age and sex differences in cerebral hemodynamics: a transcranial Doppler study. Stroke, 29(5), 963–967. 10.1161/01.str.29.5.963

Mayeux, R., Stern, Y., Rosen, J., & Leventhal, J. (1981). Depression, intellectual impairment, and Parkinson disease. Neurology, 31(6), 645–650. 10.1212/wnl.31.6.645

McDonough, I. M., Bischof, G. N., Kennedy, K. M., Rodrigue, K. M., Farrell, M. E., & Park, D. C. (2016). Discrepancies between fluid and crystallized ability in healthy adults: a behavioral marker of preclinical Alzheimer’s disease. Neurobiology of Aging, 46, 68–75. 10.1016/j.neurobiolaging.2016.06.011

Muggeo, V. M. R. (2016). Testing with a nuisance parameter present only under the alternative: a score-based approach with application to segmented modelling. Journal of Statistical Computation and Simulation, 86(15), 3059–3067. 10.1080/00949655.2016.1149855

Nasreddine, Z. S., Phillips, N. A., Bédirian, V., Charbonneau, S., Whitehead, V., Collin, I., Cummings, J. L., & Chertkow, H. (2005). The Montreal Cognitive Assessment, MoCA: a brief screening tool for mild cognitive impairment. Journal of the American Geriatrics Society, 53(4), 695–699. 10.1111/j.1532-5415.2005.53221.x

Pantoni, L., & Garcia, J. H. (1997). Pathogenesis of leukoaraiosis: a review. Stroke, 28(3), 652–659. 10.1161/01.str.28.3.652

Pantoni, L., Garcia, J. H., & Gutierrez, J. A. (1996). Cerebral white matter is highly vulnerable to ischemia. Stroke, 27(9), 1641–1646; discussion 1647. 10.1161/01.str.27.9.1641

Pearson, J. D., Morrell, C. H., Brant, L. J., Landis, P. K., & Fleg, J. L. (1997). Age-associated changes in blood pressure in a longitudinal study of healthy men and women. *The Journals of Gerontology. Series A*, Biological Sciences and Medical Sciences, 52(3), M177–83. 10.1093/gerona/52a.3.m177

Penotti, M., Farina, M., Sironi, L., Barletta, L., Gabrielli, L., & Vignali, M. (1996). Cerebral artery blood flow in relation to age and menopausal status. Obstetrics and Gynecology, 88(1), 106–109. 10.1016/0029-7844(96)00119-6

Preacher, K. J., & Kelley, K. (2011). Effect size measures for mediation models: Quantitative strategies for communicating indirect effects. In Psychological Methods (Vol. 16, pp. 93–115). American Psychological Association. 10.1037/a0022658

Purkayastha, S., & Sorond, F. (2012). Transcranial Doppler ultrasound: technique and application. Seminars in Neurology, 32(4), 411–420. 10.1055/s-0032-1331812

Reis, S. E., Gloth, S. T., Blumenthal, R. S., Resar, J. R., Zacur, H. A., Gerstenblith, G., & Brinker, J. A. (1994). Ethinyl estradiol acutely attenuates abnormal coronary vasomotor responses to acetylcholine in postmenopausal women. Circulation, 89(1), 52–60. 10.1161/01.cir.89.1.52

Reslan, O. M., & Khalil, R. A. (2012). Vascular effects of estrogenic menopausal hormone therapy. Reviews on Recent Clinical Trials, 7(1), 47–70. 10.2174/157488712799363253

Rosseel, Y. (2012). lavaan: An R Package for Structural Equation Modeling. Journal of Statistical Software, *48*(2 SE-Articles), 1–36. 10.18637/jss.v048.i02

RStudio Team. (2020). RStudio: Integrated Development for R. http://www.rstudio.com/

Salthouse, T. (2012). Consequences of age-related cognitive declines. Annual Review of Psychology, 63, 201–226. 10.1146/annurev-psych-120710-100328

Schneider, J. A., Arvanitakis, Z., Bang, W., & Bennett, D. A. (2007). Mixed brain pathologies account for most dementia cases in community-dwelling older persons. Neurology, 69(24), 2197–2204. 10.1212/01.wnl.0000271090.28148.24

Skurnick, J. H., Aladjem, M., & Aviv, A. (2010). Sex differences in pulse pressure trends with age are cross-cultural. Hypertension (Dallas, Tex. : 1979), 55(1), 40–47. 10.1161/HYPERTENSIONAHA.109.139477

Staessen, J. A., Gasowski, J., Wang, J. G., Thijs, L., Den Hond, E., Boissel, J. P., Coope, J., Ekbom, T., Gueyffier, F., Liu, L., Kerlikowske, K., Pocock, S., & Fagard, R. H. (2000). Risks of untreated and treated isolated systolic hypertension in the elderly: meta-analysis of outcome trials. *Lancet (London*, England), 355(9207), 865–872. 10.1016/s0140-6736(99)07330-4

Stamler, J., & Neaton, J. D. (2008). The Multiple Risk Factor Intervention Trial (MRFIT)--importance then and now. JAMA, 300(11), 1343–1345. 10.1001/jama.300.11.1343

Steppan, J., Barodka, V., Berkowitz, D. E., & Nyhan, D. (2011). Vascular stiffness and increased pulse pressure in the aging cardiovascular system. Cardiology Research and Practice, 2011, 263585. 10.4061/2011/263585

Tan, C. H., Low, K. A., Chiarelli, A. M., Fletcher, M. A., Navarra, R., Burzynska, A. Z., Kong, T. S., Zimmerman, B., Maclin, E. L., Sutton, B. P., Gratton, G., & Fabiani, M. (2019). Optical measures of cerebral arterial stiffness are associated with white matter signal abnormalities and cognitive performance in normal aging. Neurobiology of Aging, 84, 200–207. 10.1016/j.neurobiolaging.2019.08.004

Tan, C. H., Low, K. A., Kong, T., Fletcher, M. A., Zimmerman, B., Maclin, E. L., Chiarelli, A. M., Gratton, G., & Fabiani, M. (2017). Mapping cerebral pulse pressure and arterial compliance over the adult lifespan with optical imaging. PloS One, 12(2), e0171305. 10.1371/journal.pone.0171305

Tarantini, S., Tran, C. H. T., Gordon, G. R., Ungvari, Z., & Csiszar, A. (2017). Impaired neurovascular coupling in aging and Alzheimer’s disease: Contribution of astrocyte dysfunction and endothelial impairment to cognitive decline. Experimental Gerontology, 94, 52–58. 10.1016/j.exger.2016.11.004

Team, R. C. (2022). R: A language and environment for statistical computing. R Foundation for Statistical Computing. https://www.r-project.org/

Unsworth, N., Heitz, R. P., & Engle, R. W. (2005). An automated version of the operation span task. Behavior Research Methods, 37(3), 498–505.

Villeneuve, S., & Jagust, W. J. (2015). Imaging Vascular Disease and Amyloid in the Aging Brain: Implications for Treatment. The Journal of Prevention of Alzheimer’s Disease, 2(1), 64–70. 10.14283/jpad.2015.47

Vito, R., & Muggeo, M. (2008). segmented: An R Package to Fit Regression Models with Broken-Line.

Wei, K., Tran, T., Chu, K., Borzage, M. T., Braskie, M. N., Harrington, M. G., & King, K. S. (2019). White matter hypointensities and hyperintensities have equivalent correlations with age and CSF β-amyloid in the nondemented elderly. Brain and Behavior, 9(12), e01457. 10.1002/brb3.1457

Wetter, N. C., Hubbard, E. A., Motl, R. W., & Sutton, B. P. (2016). Fully automated open-source lesion mapping of T2-FLAIR images with FSL correlates with clinical disability in MS. Brain and Behavior, 6(3), e00440. 10.1002/brb3.440

Xing, D., Nozell, S., Chen, Y.-F., Hage, F., & Oparil, S. (2009). Estrogen and mechanisms of vascular protection. *Arteriosclerosis*, Thrombosis, and Vascular Biology, 29(3), 289–295. 10.1161/ATVBAHA.108.182279

Zimmerman, B., Rypma, B., Gratton, G., & Fabiani, M. (2021). Age-related changes in cerebrovascular health and their effects on neural function and cognition: A comprehensive review. Psychophysiology, 58(7), e13796. 10.1111/psyp.13796

