## Supplementary Material for "Neurovascular Mechanisms of Cognitive Aging: Sex-Related Differences in the Average Progression of Arteriosclerosis, White Matter Atrophy, and Cognitive Decline"

**Supplementary Analyses**

**Statistical Harmonization Procedure—Details**

**Detailed steps of the statistical harmonization procedure employed on neuropsychological test scores are written below.**

- Step 1: Decide which theoretical constructs you are interested in (i.e., crystallized cognition, fluid cognition).
- Step 2: For each data set (i.e., study), determine which measures are thought to be useful to estimate the theoretical construct (e.g., fluid cognition: OSPAN scores, Raven’s progressive matrices, Wisconsin Card Sorting Test, Pattern Comparison Processing Speed—NIH; crystallized cognition: Shipley’s vocabulary test, KBIT-2 verbal scores, Oral Reading Recognition—NIH).
- Step 3: For each individual variable, compute standardized scores within a data set (i.e., study). If multiple variables are used for a particular psychological construct, average the values together, and then standardize the scores again.
- Step 4: Determine an age group for which all data set overlap (in this case, 50-70 years old). These sub-samples are now assumed to be equivalent across different data sets — meaning that they have the same mean and standard deviation. The size of each sub-sample by self-reported sex is listed below:

**Study 1**: *n_Female_* = 21; *n_Male_* = 23

**Study 2**: *n_Female_* = 23; *n_Male_* = 21

**Study 3**: *n_Female_* = 44; *n_Male_* = 22

**Study 4**: *n_Female_* = 29; *n_Male_* = 20

- Step 5: Compute the means and standard deviations of each construct for these sub-samples, separately for each subsample, and use them to “re-standardize” the data of different sub-samples (meaning, for each data set, subtract from all individuals the mean of the subsample belonging to that dataset and divide by the standard deviation of the sub-sample of that dataset). At this point, across all data sets, the means of data falling within the age range of the sub-sample will be 0, and the standard deviation 1. But the data not falling within this range will have different values. This operation should be repeated for each construct.
- Step 6: To make the scale for different constructs comparable to each other, standardize each of them again after having pooled together the different data sets.

**Statistical Harmonization Procedure—Validation**

The cognitive aging literature has repeatedly demonstrated, using both cross-sectional and longitudinal data, differential age-related trends for fluid versus crystallized abilities. Specifically, for fluid cognition, a *linear* age-related decline is commonly observed, whereas for crystallized (verbal) cognition, a *non-linear* age-related trend is generally observed such that crystallized abilities increase throughout the lifespan until approximately age 70, but decrease precipitously afterwards **(e.g.,** **see Salthouse, 2012, Figure 2 for plots illustrating these general trends)**.

To provide supporting evidence for the validity of the statistical harmonization procedure employed, we conducted separate second-order polynomial regressions with *age* (linear term) and *age^2^* (quadratic term) as the two predictors and composite fluid score or composite crystallized score as the outcome. If our harmonization procedure is valid, then the linear term should be a stronger predictor and account for more variance than the quadratic term in the fluid ability regression, thereby replicating the results of past aging research. However, for the crystallized ability regression, quadratic term should be a stronger predictor and account for more variance than the linear term, emulating previous studies.

As multicollinearity, or high correlation among powers of the explanatory variable, is a common problem in polynomial regression, the polynomial terms were orthogonalized so as to be uncorrelated, using the ‘poly()’ function in R. This allowed us to better partition and assess the unique contributions of linear and quadratic terms on the outcome.

The fitted regression equations with standardized coefficients are:

a) Fluid Ability = .02 - .57*Age - .18*Age^2^
b) Crystallized Ability = -.01 + .18*Age - .19*Age^2^

First, overall, regression model ‘a’ was statistically significant (*R*^2^ = .35, *F*(2, 191) = 51.45, *p* < .001). More importantly, while the quadratic term was a significant predictor after orthogonalization (*R_p_*^2^= .05 ,β = -.18, *p* < .001), the magnitude of the linear term’s coefficient was larger and accounted for a higher proportion of the variance in fluid ability (*R_p_*^2^= .33 ,β = -.57, p < .001) than the quadratic term (as indicated by partial R-squared: .05 versus .33), indicating that the relation between age and our harmonized fluid ability scores is better described by a negative linear trend, which is consistent with the literature.

Next, overall, regression model ‘b’ was statistically significant (*R*^2^ = .07, *F*(2, 191) = 6.83, *p* < .001). More importantly, while the linear term was a significant predictor after orthogonalization (*R_p_*^2^= .03, β = .18, *p* = .01), the quadratic term was a slightly stronger predictor of crystallized ability (*R_p_*^2^= .04, β = -.19, *p* = .01), indicating that the relation between age and our harmonized crystallized ability scores is possibly better described by a nonlinear trend, which is also consistent with the literature.

Scatterplots of the relationship between age and fluid or crystallized abilities, with linear and quadratic trends superimposed, are presented below.

**Figure S1**

Scatterplot of the relationship between age and composite fluid ability scores, post-harmonization. Participants are color-coded by the study they participated in. The red line/trend indicates the predictor (here, the *linear* term) that is a stronger predictor and accounts for a larger proportion of the variance in the outcome, fluid ability.

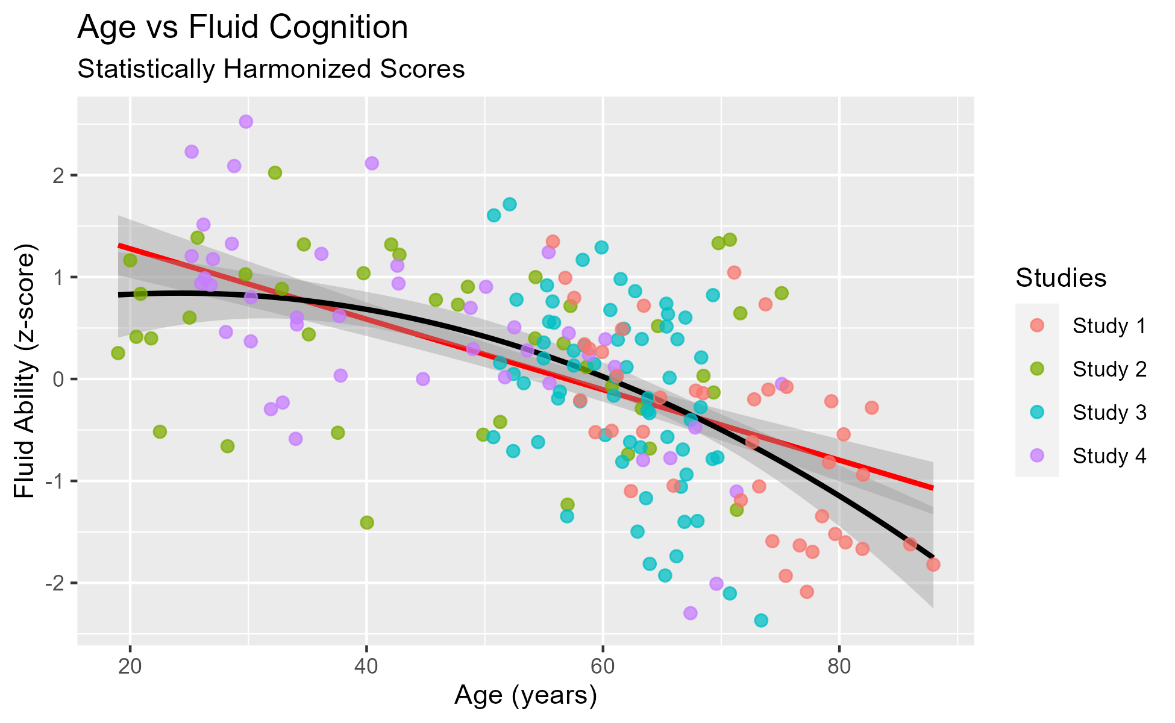

**Figure S2**

Scatterplot of the relationship between age and composite crystallized ability scores, post-harmonization. Participants are color-coded by the study they participated in. The red line/trend indicates the predictor (here, the *quadratic* term) that is a slightly stronger predictor and accounts for a larger proportion of the variance in the outcome, crystallized ability.

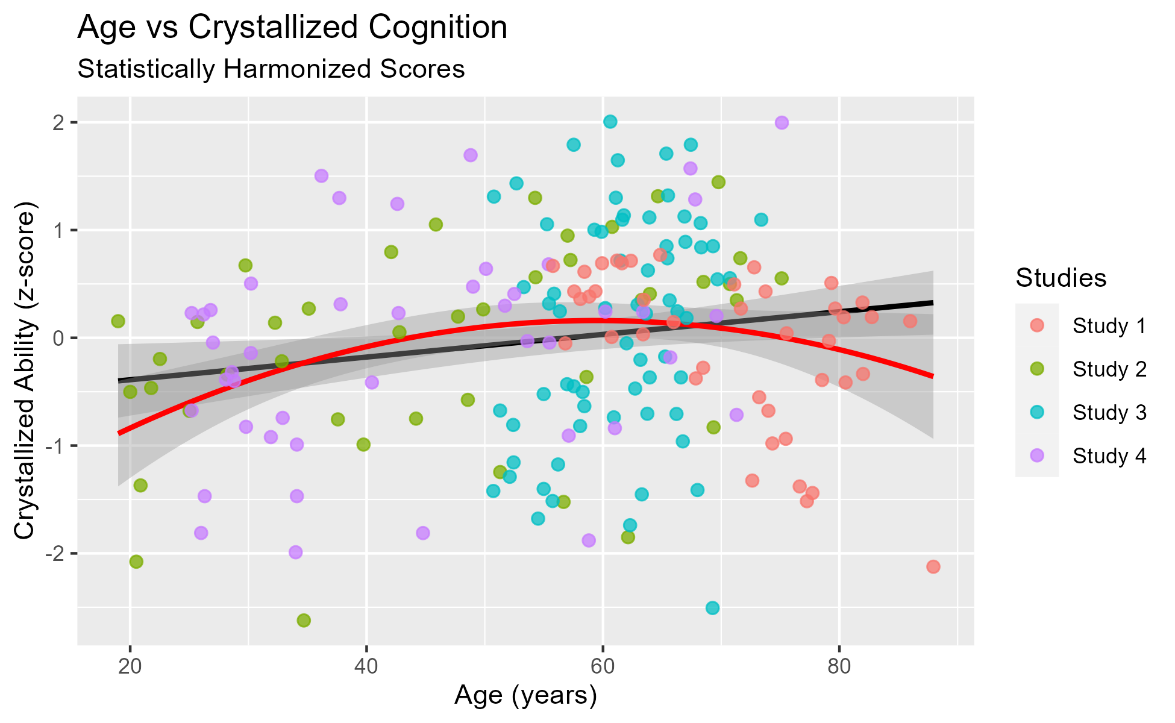

**Piecewise Regression Analyses (Systemic Pulse Pressure)**

We repeated the piecewise regression analyses detailed in the main text with age and systemic pulse pressure, instead of PReFx, in an effort to replicate the optical cerebral findings. Below, we plot bilinear trends for all participants. In a separate plot, we show the bilinear patterns by self-reported sex. The analyses reveal breakpoints that are highly similar to those observed in the main text with PReFx. Overall, a statistically breakpoint is observed (age = 51.29 years, *SE* = 3.58, *p* < .001; 95% CI [44.25, 58.34]), with only the post-breakpoint slope significant, indicating an acceleration in arterial stiffening after this age.

The plot demonstrating sex differences in the age-related progression of arteriosclerosis mimics that of PReFx. That is, only females exhibit a significant breakpoint (age = 50.10 years, *SE* = 3.57, *p* < .001; 95% CI [43.03, 57.17]), indicating an acceleration in arterial stiffening, presumably due to menopause-induced decreases in estrogen, resulting in a diminution of anti-inflammatory and vasoprotective mechanisms (Xing et al., 2009).

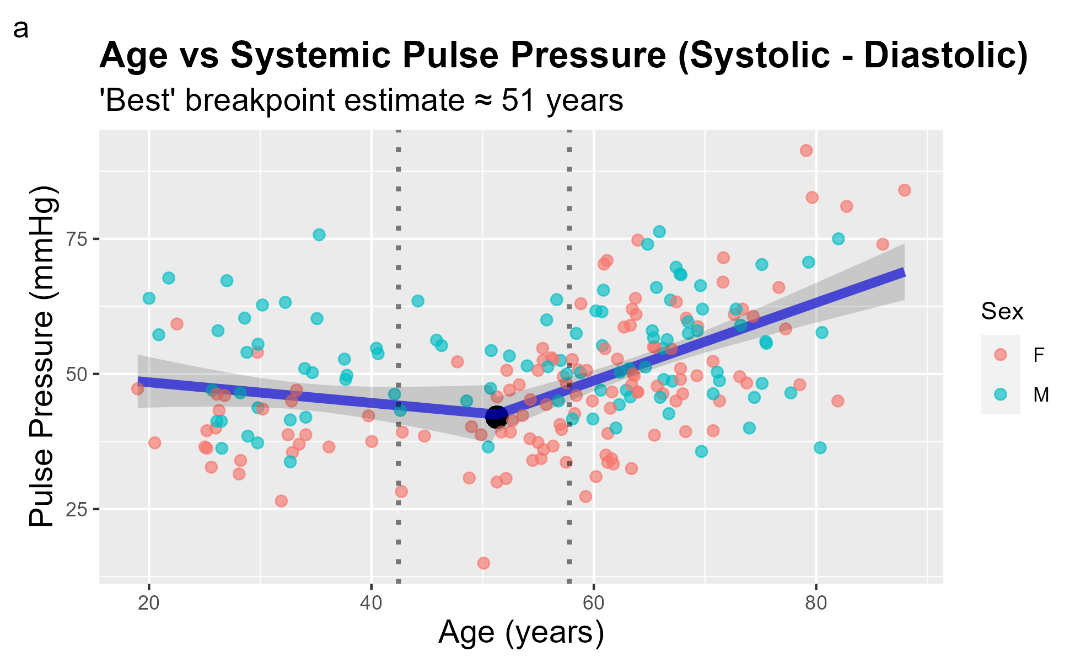

**Figure S3**

Scatterplot of the relationship between age and pulse pressure. Participants are color-coded by their self-reported sex. A significant breakpoint was estimated at approximately 51 years, suggesting a general acceleration in arteriosclerosis progression.

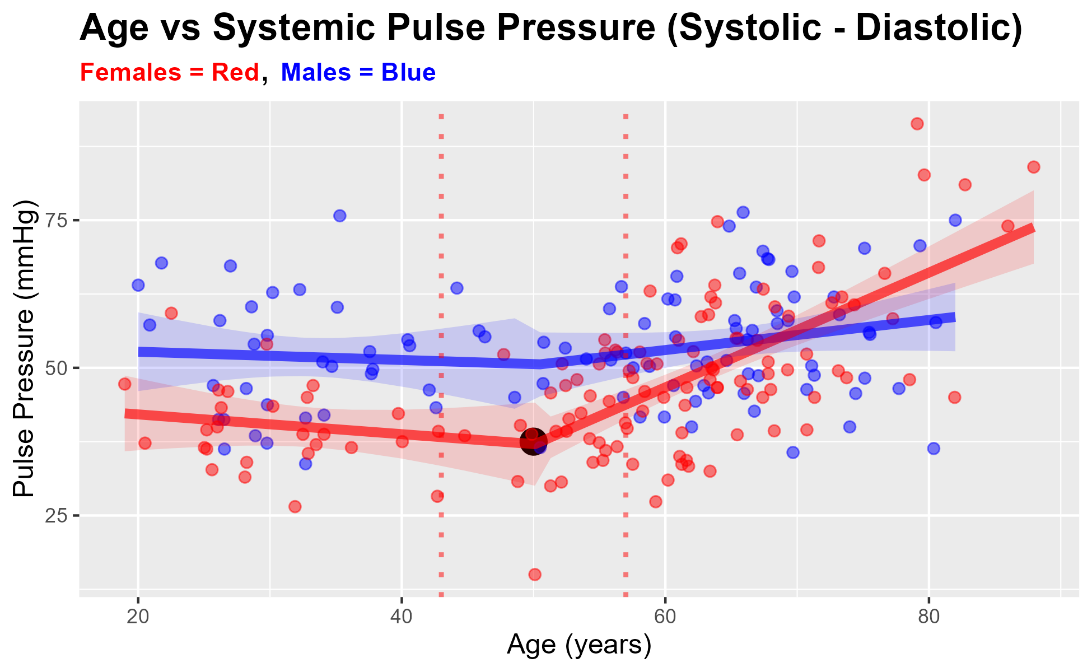

**Figure S4**

Scatterplot of the relationship between age and pulse pressure by sex. Participants are color-coded by their self-reported sex. Red and blue shaded areas indicate the 95% confidence interval bands for females and males, respectively. Only females exhibited a significant breakpoint, occurring at approximately age 50. This suggests that females are more susceptible to vascular injury later in life, likely due to the decreased production of estrogen, a consequence of menopause.

**Two-Mediator Serial/Sequential Mediation (Systemic Pulse Pressure)**

In an attempt to replicate the effects of age-related declines in measures of cerebrovascular health and white matter integrity on fluid and crystallized cognition, we conducted additional two-mediator serial mediation analyses with systemic pulse pressure (instead of PReFx) as the index of vascular health. As illustrated in the main text via two mediation analyses, when PReFx (an index of cerebral arterial health) served as the proxy for vascular health, both analyses provided evidence consistent with the cascade model of neurocognitive aging. That is, for both analyses, we observed a statistically significant *multiple-mediator* (serial) indirect effect (tested via p-values generated from bootstrapped distributions of 5,000 samples) such that age-related declines in vascular health seemingly indirectly affect fluid abilities via their deleterious effect on white matter integrity (i.e., white matter lesions), while crystallized abilities were left relatively unscathed. Self-reported biological sex is controlled for in the regression-based mediation analyses.

The aforementioned analyses were repeated with systemic pulse pressure as the vascular index, while also controlling for sex. Results are illustrated below via path diagrams with standardized path coefficients and bootstrapped p-values based on 5,000 bootstrap samples (Rosseel, 2012). The pattern of significant results replicates those observed in the main text.

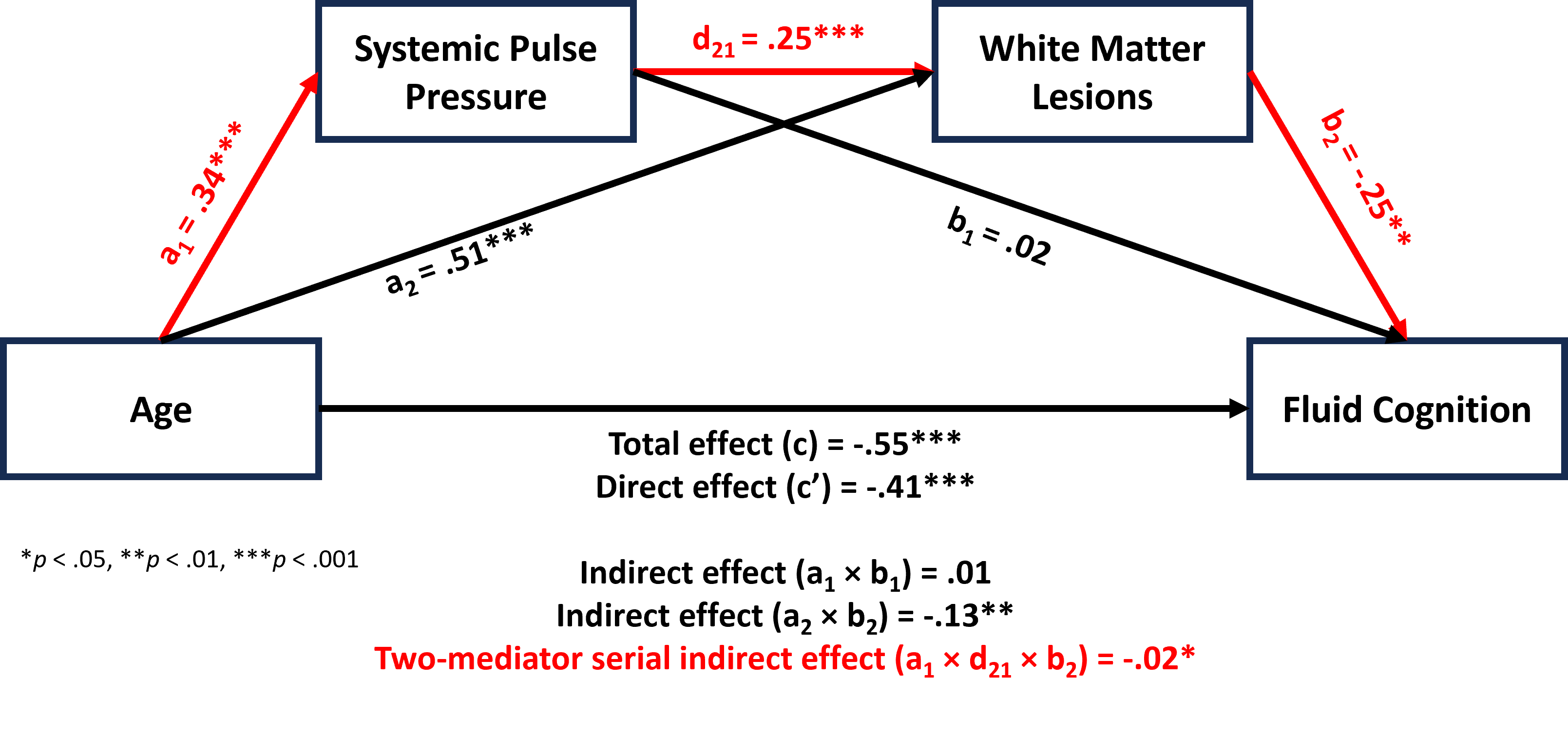

**Figure S5**

The presence of a significant two-mediator sequential indirect effect (illustrated in red text) suggests that age-related declines in systemic vascular health and white matter integrity may significantly impact fluid abilities throughout the lifespan. Self-reported sex is controlled for in these analyses but is not included in the diagram to increase visual clarity.

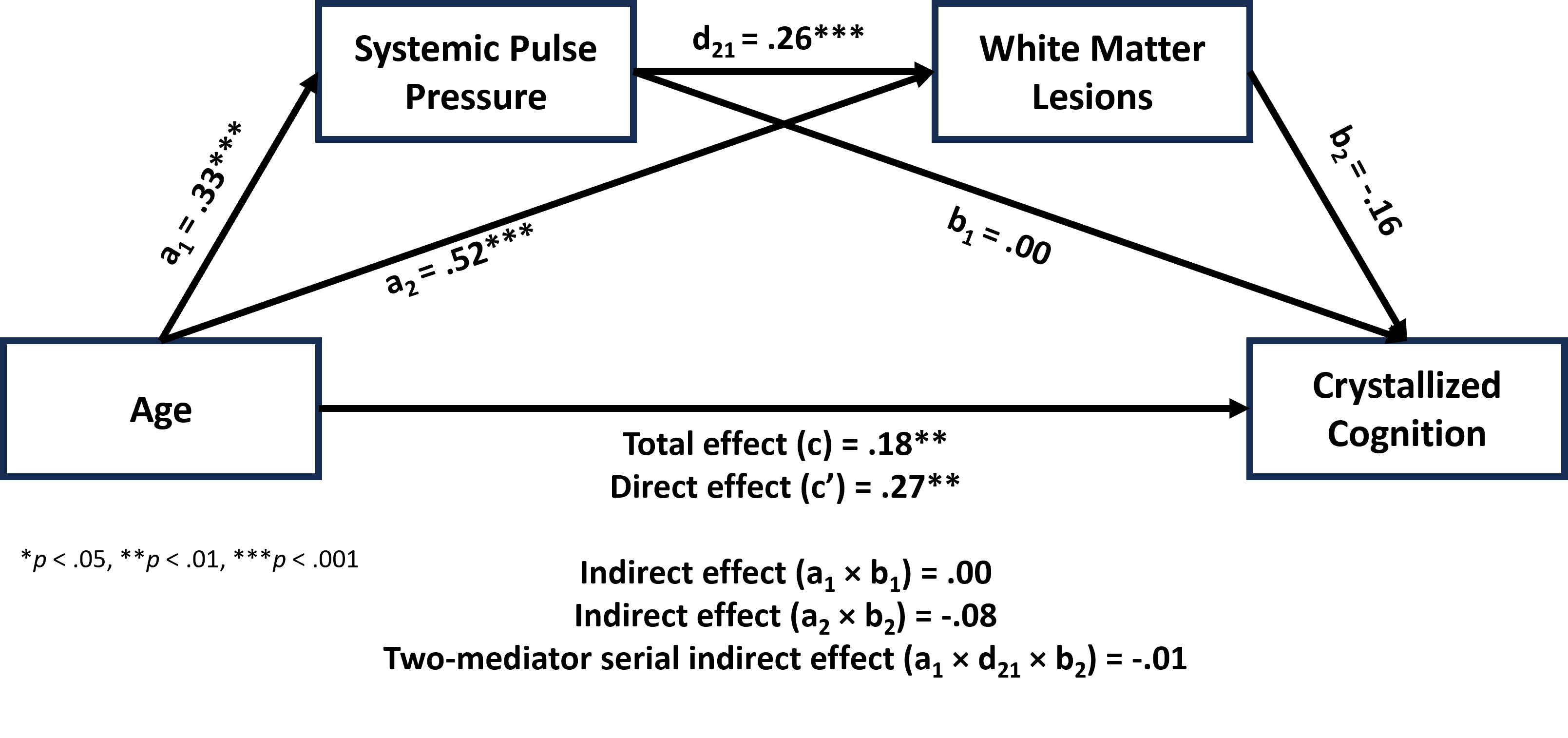

**Figure S6**

Given the absence of a significant two-mediator indirect effect, these analyses suggest that age-related declines in systemic vascular health and white matter integrity may not significantly impact crystallized abilities throughout the lifespan. Self-reported sex is controlled for in these analyses but is not included in the diagram to increase visual clarity.

**Manually Corrected vs Uncorrected FreeSurfer WMH Volumes**

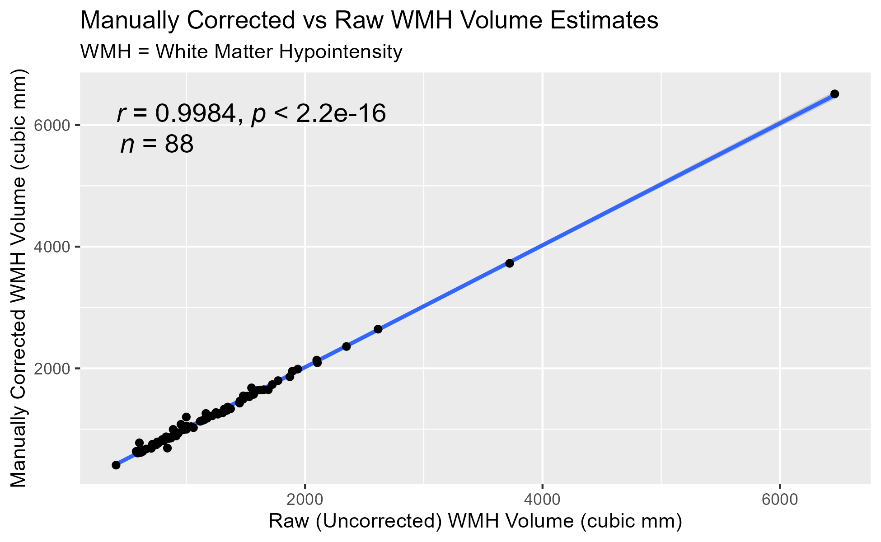
As we only have manually corrected FreeSurfer volume, thickness, and surface area estimates for a subset of subjects (and because it can take considerable time to inspect each participant’s MPRAGE manually and thoroughly for segmentation errors), in order to expand our sample size, we needed to confirm that the corrected and uncorrected white matter hypointensity (WMH) volume estimates were highly correlated. Therefore, in a subset of subjects who had both measures (*n* = 88), we examined the linear correlation between manually corrected and raw FreeSurfer WMH volumetric estimates. The results, illustrated via **Figure S7** below, are extremely reassuring, as the correlation between the two estimates is essentially equal to 1, indicating a near-perfect correlation. Therefore, for all analyses, we utilize the raw FreeSurfer estimates of white matter hypointensity volume.

**Figure S7**

**Comparison of T1-Weighted White Matter Hypointensity and T2-FLAIR White Matter Hyperintensity Volumes**

Extant evidence suggests that T1w WM-hypo volumes and T2-FLAIR WM-hyper volumes assess largely overlapping cerebral white matter phenomena during aging, including non-specific lesions, which may manifest as axonal demyelination, axonal degeneration/loss, and reactive gliosis (Pantoni & Garcia, 1997). Among 56 non-demented elderly individuals, Wei et al. (2019) demonstrated that WM-hypointensity volumes and WM-hyperintensity volumes were highly correlated (*r* = .81). Furthermore, they showed that hypointensities and hyperintensities exhibited highly similar correlations with other variables of interest, including age and CSF biomarkers of Alzheimer’s disease, including β-amyloid and tau deposition. While T2-FLAIR hyperintensities may be more sensitive to periventricular and deep white matter lesions, this study strongly suggests that T1w white matter hypointensities can serve as an adequate proxy for cerebral white matter lesions, in the absence of T2-FLAIR imaging. Accordingly, as a subsample of our participants underwent both T1-weighted and T2-FLAIR imaging, we attempted to replicate the findings of Wei and colleagues (2019), thereby assuaging concerns regarding our employment of T1w white matter hypointensity volume as an index of overall white matter lesion burden. Below, utilizing a limited subsample of participants (*N* = 47; **Figure S8**), we show that WM-hypo and WM-hyper volumes are strongly correlated (*r* = .70).

**T2-FLAIR Lesion Mapping Method**

A subset of participants in studies 3 and 4 underwent T2-weighted fluid-attenuated inversion recovery (FLAIR) imaging in a 3-Tesla Siemens Prisma MR scanner, using a 20-channel head coil. The T2-FLAIR sequence utilized the following pulse parameters: TR = 6,000 ms; TE = 388 ms; TI = 2,200 ms; 208 sagittal slices; slice thickness = 1.00 mm; voxel sizes = 1.0 × 1.0 × 1.0 mm, acceleration factor = 2.

Lesion-mapping was performed using an unsupervised, single-image method described in Wetter et al., 2016, based on tools in the FMRIB Software Library (FSL) (Smith et al., 2004). First the FLAIR image was bias-field corrected to correct non-uniformity in the low frequency intensity using a tool based on FSL's brain extraction tool (BET) (Smith 2002). The FLAIR image was skull stripped in a two-step process. The FreeSurfer brain mask was coregistered into subject space and used to extract the brain. Usually, some voxels of dura still exist in this brain extraction. Therefore, the second step used BET to refine the brain extraction from FreeSurfer. Then Gaussian fitting of the intensity histogram was used by FMRIBs Automated Segmentation Tool (FAST) to segment brain images by tissue type (Zhang et al., 2001). Voxels comprising the hyperintensities appear as outliers of the fitted Gaussian distribution of brain tissue on the intensity histogram. The darkest bins in the histogram are iteratively removed until an empty bin is found, separating the histogram of normal tissue intensities from the hyperintensities. This bin value was used as a threshold to remove the brain tissue.

Although this method captures the hyperintense lesions, it erroneously labels the septum pellucidum, small regions in the peripheral gray matter, and sometimes the midbrain gray matter. In order to clean the images, Wetter and colleagues used a FLAIR-like standard space, which was created by subtracting the ICBM CSF mask from the ICBM T2 standard mask. Using their strategy, we utilized FLIRT (Jenkinson and Smith, 2001; Jenkinson et al., 2002) and FNIRT (Smith 2007) to warp the FLAIR-like standard image to subject space, and used the transformation to bring the ICBM WM mask to the subject space. Voxels with a white matter probability above 0.6 were included in the mask. Lesions were selected only if they included at least one voxel within the WM mask, in order to eliminate false positives outside of the WM. Midline false positives were removed by eliminating any lesions that were within 4mm of the mid-sagittal plane.

To identify and quantify white matter lesion load, we developed on top of existing automated lesion-mapping code originally developed and described in (Wetter et al., 2016). The new developments were made to reduce the false positive identification of lesions and incorporate the ability to separate periventricular from deep white matter lesions.

The transformation of the ICBM WM mask into subject space sometimes resulted in some of the gray matter structures in subject space overlapping with the white matter mask. This particularly posed a problem when the white matter mask did not sufficiently exclude the caudate and ventricles, where it can be difficult to separate periventricular lesions from false positive voxels. To further ensure that this did not happen, we used the FreeSurfer segmentations of the ventricles and caudate as additional masks to exclude hyperintense voxels.

Finally, periventricular lesions were separated from deep white matter lesions. This is done using an iterative procedure. First, the hyperintensities bordering the ventricles are identified using the later ventricle mask from FreeSurfer. Then hyperintensities touching these voxels are identified and added to the growing periventricular lesion mask. This continues until there are no more neighboring hyperintensities, marking the end of any contiguous lesions radiating out from near the ventricles. Since the rest of the hyperintensities have been masked so as to appear only in the other white matter, all non-periventricular lesions left in the lesion mask are classified as deep white matter lesions. The final lesion masks were then manually inspected using quality control images output by the developed method.

We also upgraded the usability and shareability of the lesion-mapping code to make it much easier to use as a community tool (https://github.com/mrfil/lesion-mapper-bids/). We did this by rewriting the code to accept Brain Imaging Data Structure (BIDS) datasets, which is quickly becoming the standard organizational structural for brain imaging data and incorporating a variety of parameters that help adjust the sensitivity or specificity of the lesion mapping. The code is actively developed and shared on github, so that anyone with a BIDS dataset with FLAIR data is able to run the automated lesion mapping software with a single command. The method includes options to leverage good anatomical segmentations from fmriprep or FreeSurfer, commonly used tools in the MRI community, if they exist, in order to improve the lesion map.

Research has shown that the mechanisms underlying periventricular white matter lesion formation may differ from deep white matter lesion formation (Nyquist et al., 2015; Alber et al., 2019; Dadar et al., 2019; Min et al., 2021). Evidence of the two types of lesions may be predictive of different of subsequent changes and relate to different aspects of physiology, but this question has not been explored in detail, especially with a specific focus on the 50-70 year old age-range. This range is particularly important, as it marks the beginning of the transition from more primary aging processes to the various types of pathologies characteristic of secondary aging, including age-related cerebrovascular dysfunction. In our subsample, a striking finding is that age seems to especially predict the periventricular lesion load. There is a significant correlation between age and the log-transformed periventricular lesion volumes *r*(45) = .30, *p* = .04,which becomes non-significant when all lesions are included *r*(45) = .16, *p* = .30, demonstrating the utility of separating periventricular lesions. However, for present purposes, we include all lesions, irrespective of location/type.

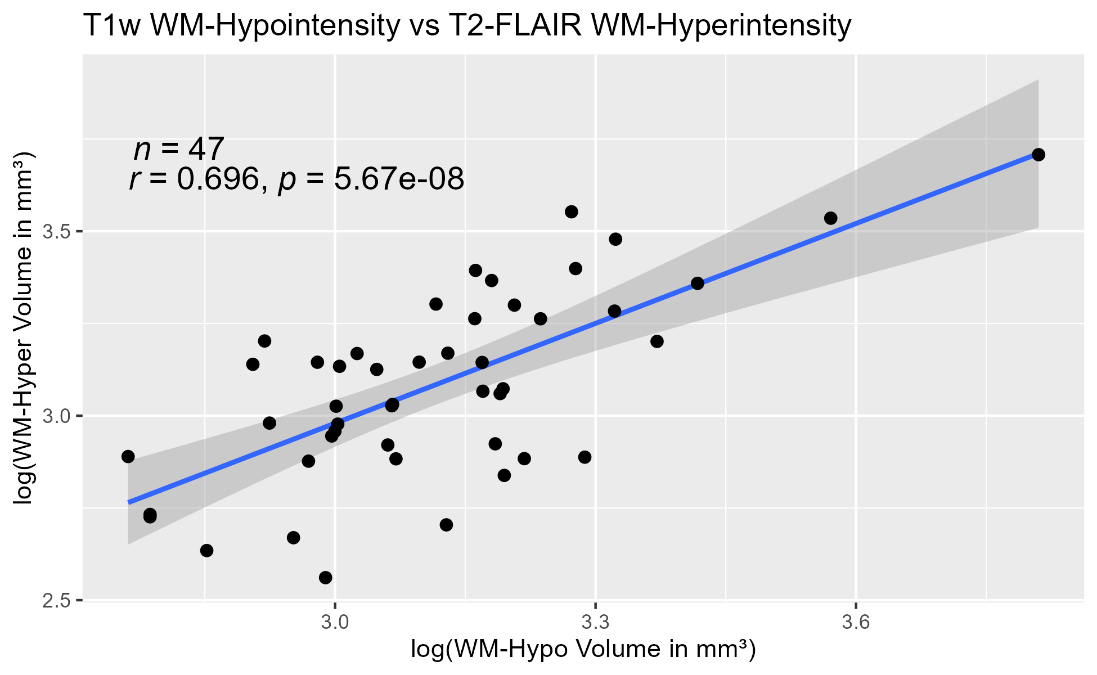

**Figure S8**

**References**

Pantoni, L., & Garcia, J. H. (1997). Pathogenesis of leukoaraiosis: a review. *Stroke*, *28*(3), 652–659. https://doi.org/10.1161/01.str.28.3.652

Rosseel, Y. (2012). lavaan: An R Package for Structural Equation Modeling. *Journal of Statistical Software*, *48*(2 SE-Articles), 1–36. https://doi.org/10.18637/jss.v048.i02

Salthouse, T. (2012). Consequences of age-related cognitive declines. *Annual Review of Psychology*, *63*, 201–226. https://doi.org/10.1146/annurev-psych-120710-100328

Wei, K., Tran, T., Chu, K., Borzage, M. T., Braskie, M. N., Harrington, M. G., & King, K. S. (2019). White matter hypointensities and hyperintensities have equivalent correlations with age and CSF β-amyloid in the nondemented elderly. *Brain and Behavior*, *9*(12), e01457. https://doi.org/10.1002/brb3.1457

Xing, D., Nozell, S., Chen, Y.-F., Hage, F., & Oparil, S. (2009). Estrogen and mechanisms of vascular protection. *Arteriosclerosis, Thrombosis, and Vascular Biology*, *29*(3), 289–295. https://doi.org/10.1161/ATVBAHA.108.182279
